# Antibiotic-dependent relationships between nasal microbiome and secreted proteome in chronic rhinosinusitis and nasal polyps

**DOI:** 10.1101/2020.04.26.060764

**Authors:** Yi-Sook Kim, Dohyun Han, Ji-Hun Mo, Yong-Min Kim, Dae Woo Kim, Hyo-Guen Choi, Jong-Wan Park, Hyun-Woo Shin

**Affiliations:** Obstructive Upper airway Research (OUaR) Laboratory, Department of Pharmacology, Seoul National University College of Medicine, Seoul, Korea; Department of Biomedical Sciences, Seoul National University Graduate School, Seoul, Korea; Proteomics core facility, Biomedical Research Institute, Seoul National University Hospital, Seoul, Korea; Department of Otorhinolaryngology-Head and Neck Surgery, Dankook University Hospital, Cheonan, Korea; Department of Otorhinolaryngology-Head and Neck Surgery, Chungnam National University Hospital, Daejeon, Korea; Department of Otorhinolaryngology-Head and Neck Surgery, Boramae Medical Center; Seoul, Korea; Department of Otorhinolaryngology-Head & Neck Surgery, Hallym University College of Medicine, Anyang, Korea; Ischemic/hypoxic Disease Institute, Seoul National University College of Medicine, Seoul, Korea; Cancer Research Institute, Seoul National University College of Medicine, Seoul, Korea; Department of Otorhinolaryngology-Head and Neck Surgery, Seoul National University Hospital, Seoul, Korea; Clinical Mucosal Immunology Study Group

**Keywords:** Metagenomics, proteomics, anti-bacterial agents, sinusitis, microbiota, proteome, mucus

## Abstract

**Background:** Antibiotics are commonly prescribed to treat chronic rhinosinusitis (CRS); however, the effects of antibiotics on the microbiome and secreted proteome remain unknown in regard to CRS.

**Objective:** We analyzed the effects of antibiotics on the nasal microbiome and secreted proteome in the context of CRS using multi-omic analysis.

**Methods:** Nasal secretions were collected from 29 control, 30 CRS patients without nasal polyps (CRSsNP), and 40 CRS patients with nasal polyps (CRSwNP). A total of 99 subjects were divided into two groups that included subjects who had taken antibiotics 3 months prior to sampling (ABX) and those who had not (NABX). We performed 16S rDNA sequence analyses and Orbitrap mass spectrometry-based proteomic analyses in data-independent acquisition (DIA) on the nasal secretions. Spearman correlation was used to assess the correlations between the nasal microbiome and secreted proteome.

**Results:** We observed a strong association between the nasal microbiome and secreted proteome according to disease status. Antibiotic use reduced differences in the microbial community and secreted proteome according to disease status. Interestingly, in nasal polyp (NP) patients, antibiotics exhibited strong effects not only on the nasal microbiome and the secreted proteome but also on their associations. Additionally, their correlations were strengthened in subjects who had taken antibiotics.

**Conclusion:** Integrative analyses revealed that the correlations between the microbiome and the secreted proteome could be altered and strengthened in subjects who used antibiotics. These findings provide novel insight into the effects of antibiotics on the nasal environment and the host responses in CRS.

## Introduction

Chronic rhinosinusitis (CRS) is a persistent inflammatory condition of the nasal mucosa and sinus. It is one of the most common upper airway diseases in Western countries and in Asia(Zhang et al. 2017). Indeed, this disease exhibits considerable heterogeneity at the clinical and molecular pathophysiologic levels(Tomassen et al. 2016). Based on this, previous studies have demonstrated that the endotypes of CRS could be characterized based on cytokines(Tomassen et al. 2016), symptoms(Divekar et al. 2015), microbiota composition(Cope et al. 2017), and clinical characteristics(Liao et al. 2018).

Recently, multi-omic analyses were performed to reveal interrelationships among the microbiome, metabolome, transcriptome, and proteome in association with human diseases(Misra et al. 2018). For example, molecular profiles of host (transcriptomics and genomics) and microbial profiles (metagenomics, metaproteomics, and metatranscriptomics) revealed that profiling of disease-associated microbiomes should be accounted for corresponding molecular changes in the host epithelium(Lloyd-Price et al. 2019). Metagenomic, transcriptomic, and proteomic analyses identified the interactions between the sputum microbiome and host interferon signaling in chronic obstructive pulmonary disease (COPD) patients(Wang et al. 2019). Therefore, integrative analyses of metagenomics and secreted proteomics in CRS could provide a better understanding of heterogeneity based on associated changes in nasal environment (microbiome) and host response (secreted proteome).

For the medical treatment of CRS, antibiotics such as penicillins/betalactams and macrolides were prescribed in about 70% of CRS visits(Smith et al. 2013). However, most of previous metagenomic analyses(Aurora et al. 2013; Choi et al. 2014; Biswas et al. 2015; Ramakrishnan et al. 2015; Biswas et al. 2017; Cope et al. 2017; Hoggard et al. 2017; Jain et al. 2017; Lal et al. 2017; Ramakrishnan et al. 2017; Chalermwatanachai et al. 2018; Copeland et al. 2018; Koeller et al. 2018; Mahdavinia et al. 2018; Biswas et al. 2019; Gan et al. 2019; Paramasivan et al. 2019; Rom et al. 2019) examining CRS excluded patients who had taken antibiotics within approximately one month prior to sampling. Here, we performed metagenomics and proteomics analyses using nasal secretions in healthy control and CRS patients. To investigate the means by which antibiotics affect both the microbiome and the secreted proteome in CRS, the subjects were divided into two groups that included the subjects who had taken antibiotics 3 months prior to sampling (ABX) and those who had not (NABX). Furthermore, we sought to gain insight into any potential interactions between the nasal microbiome and the secreted proteome.

## Results

### Differences in the microbial composition and proteome according to disease status

From our data, we identified 1,329 operational taxonomic units (OTUs) at the genus level in 29 controls, 30 CRSsNP, and 40 CRSwNP patients. To determine if the microbial composition was different according to disease status, we analyzed the alpha and beta diversity in a total of 99 subjects. Shannon and Simpson indices were significantly increased across control to CRSwNP, while Chao1 and the number of observed OTUs were not increased (Fig. 1A and Fig. S2A). There were significant differences in microbial composition in relation to disease status (Fig. 1B).

**Figure 1.**
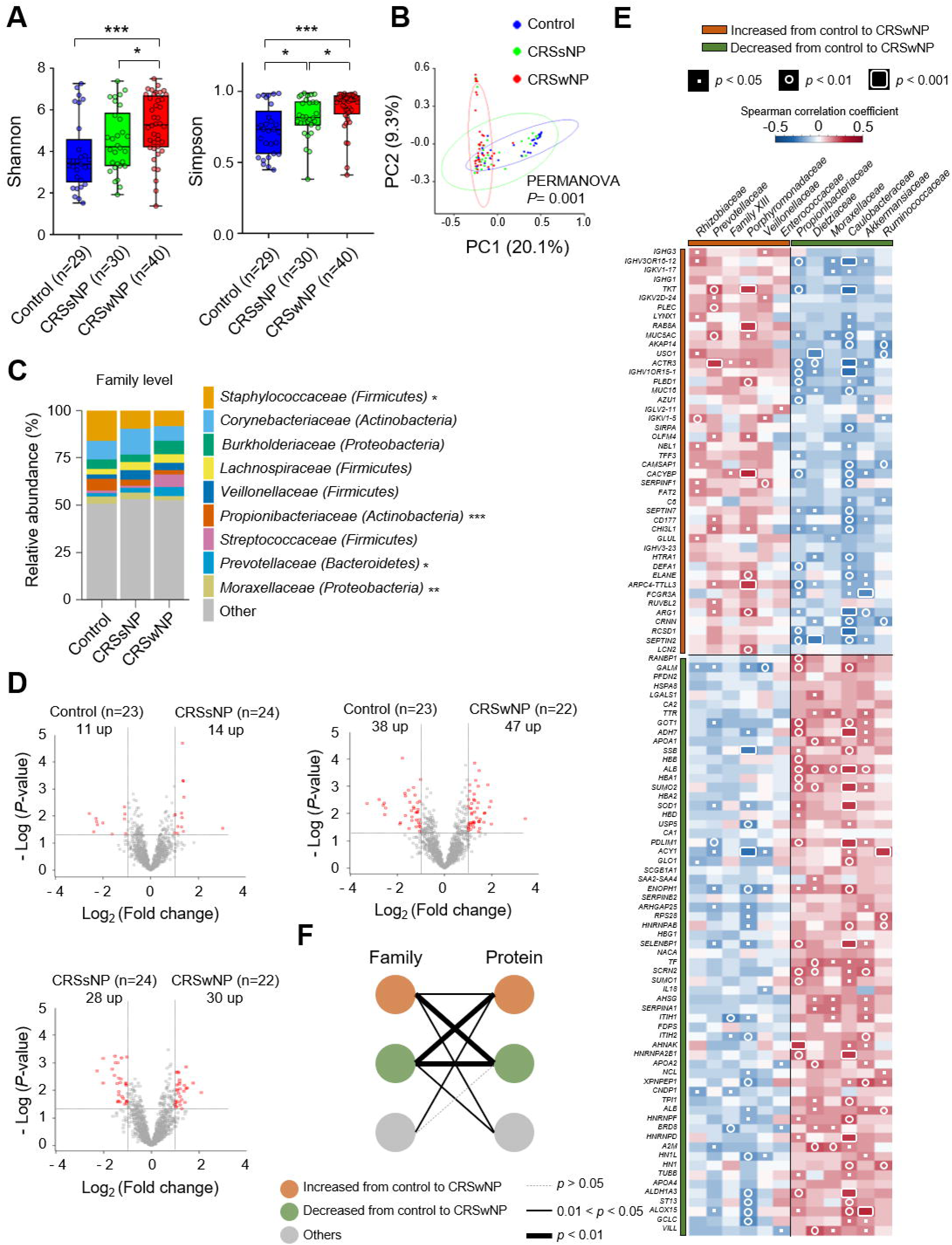
Nasal microbiome and secreted proteome profiles. (A) Alpha diversity according to disease status (horizontal line = median and whiskers = min/max range). (B) A PCoA plot based on Bray–Curtis distance matrix. (C) Distribution of bacterial families. The composition of each family exhibiting a relative abundance of greater than 3 percent is illustrated. The parenthesis represents phylum within the family. (D) Volcano plots of the proteome. The horizontal dashed line indicates *p* values of 0.05 (Student’s t-test), and the vertical dashed lines indicate fold changes of 2.0. (E) Spearman correlation heatmap of secreted proteomes (ANOVA, *p* < 0.05) and the nasal microbiome (Kruskal-Wallis, *p* < 0.05). The proteome and microbiome were arranged from top to bottom and from left to right in order of the lowest to the highest *p* value, respectively. Orange and green boxes indicated increased and decreased families and proteins from control to CRSwNP, respectively. Redundant microbiomes and proteomes were excluded from the heatmap. (F) Association between families and proteins. Orange and green circles represent the same groups previously described in (E). Gray circles represent other microbiomes and proteomes that were identified in this study. (\**p* < 0.05, \*\**p* < 0.01, \*\*\**p* < 0.001)

To investigate which bacterial taxa were different in relation to disease status, we compared the nasal bacterial composition. *Firmicutes* and *Bacteroidetes* were significantly increased from control to CRSwNP, while *Cyanobacteria* levels were significantly decreased from control to CRSwNP (Fig. S2B and C). At the family level, *Staphylococcaceae, Propionibacteriaceae*, and *Moraxellaceae* were significantly decreased from control to CRSwNP (Fig. 1C and Fig. S2D). *Prevotellaceae* was significantly increased from control to CRSwNP. Linear discriminant analysis (LDA) effect size (LEfSe) analyses revealed *Propionibacteriaceae* and *Moraxellaceae* were the most abundant in the control samples, and *Entomoplasmataceae* was the most abundant in the CRSsNP samples at the family level (Fig. S2E).

To determine if the secreted proteome was different in relation to disease status, we performed proteomic analysis of the nasal secretions of 69 subjects that were divided into 23 control, 24 CRSsNP, and 22 CRSwNP samples. We quantified a total of 2,162 proteins, and on average we quantified approximately 1,440 proteins. The number of quantified proteins from each subject is represented in Figure S3. When comparing the control and CRSsNP groups, relatively small proteomic changes were observed compared to changes observed when comparing the control and CRSwNP or CRSsNP and CRSwNP groups (Fig. 1D and Table S1).

Given that the nasal microbiome and secreted proteome were different in relation to disease status, we hypothesized that they could correlate with each other. To confirm this hypothesis, we divided the nasal microbiome into two groups that included families with increased relative abundance from control to CRSwNP (IFc) and families exhibiting decreased relative abundance from control to CRSwNP (DFc) (Kruskal-Wallis, *p* < 0.05). We also divided the secreted proteome into two groups that included proteins with increased normalized intensity from control to CRSwNP (IPc) and proteins exhibiting decreased normalized intensity from control to CRSwNP (DPc) (ANOVA, *p* < 0.05). A number of significant positive or negative correlations were observed among the groups (IFc, DFc, IPc, and DPc) (Fig. 1E). To confirm the associations among the four groups, we applied adaptive sum of powered correlation (aSPC) tests. These analyses revealed significant global association among the groups (IFc, DFc, IPc and DPc) (aSPC test, *p* < 0.01) (Fig. 1F). In contrast, less significant correlations were detected between each group (IFc, DFc, IPc and DPc) and others in regard to the microbiome and proteome, respectively (aSPC test, *p* > 0.01). These findings suggest a strong association between the nasal microbiome and the secreted proteome according to disease status.

### Differential microbiome composition according to the use of antibiotics

Further, we sought to examine variations in the microbial community in relation to disease status in NABX and ABX. The NABX group consisted of 27 control, 22 CRSsNP, and 24 CRSwNP patients, while the ABX group consisted of 2 control, 8 CRSsNP, and 16 CRSwNP patients. In NABX, unlike Chao1 and the number of observed OTUs, the Shannon and Simpson indices were significantly increased from control to CRSwNP (Fig. 2A and Fig. S4A). A clearly identifiable clustering was observed in NABX (Fig. S4C); however, in ABX, there was no significant difference in the alpha and beta diversities (Fig. 2B and Fig. S4B and D).

**Figure 2.**
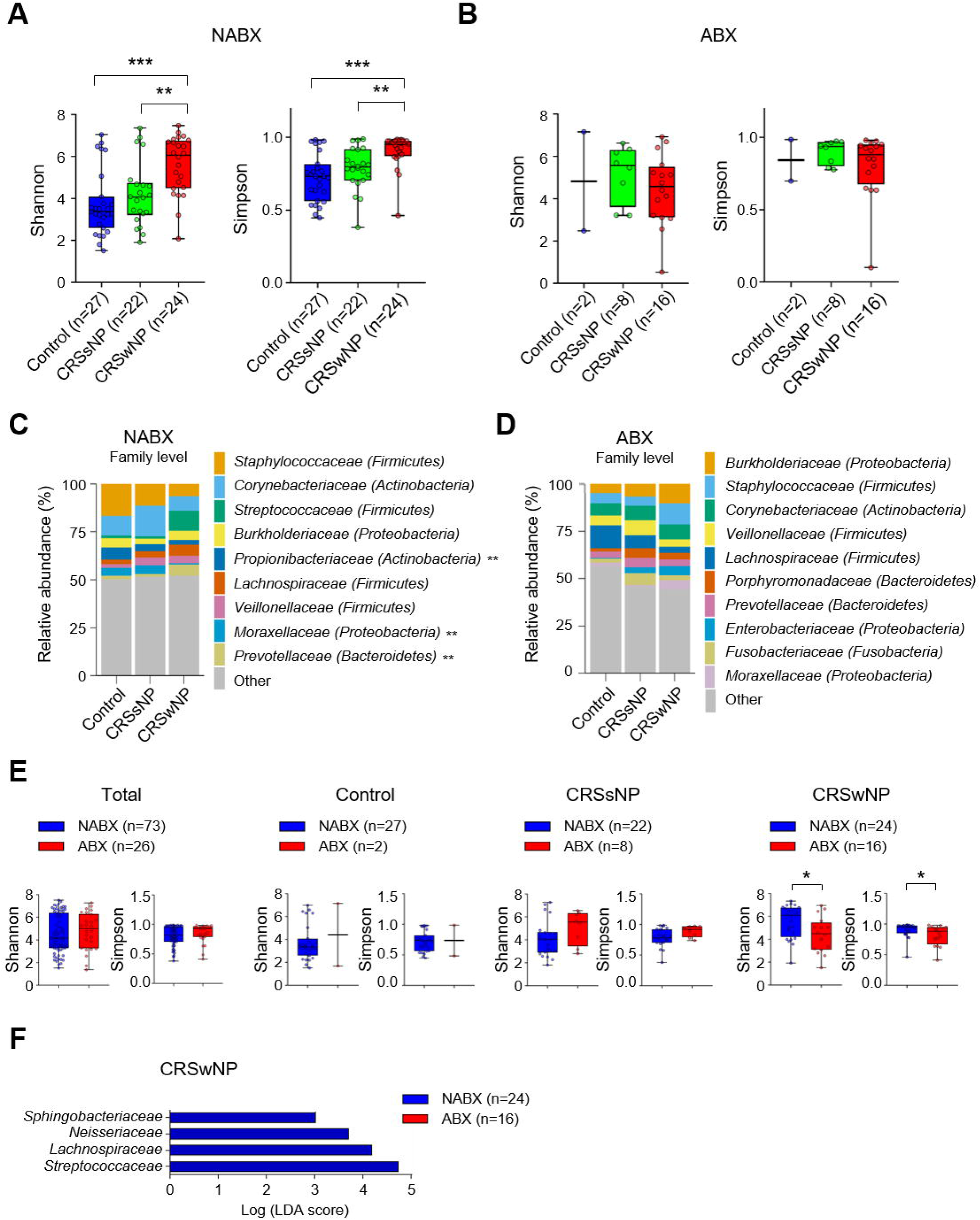
Differential microbial composition according to the use of antibiotics. (A and B) Comparison of Shannon and Simpson indices according to disease status in NABX and ABX. (C and D) Distribution of bacterial families according to disease status in NABX and ABX. The composition of each family exhibiting a relative abundance of greater than 3 percent is illustrated. The parenthesis indicates phylum within the family. (E) Comparison of Shannon and Simpson indices between NABX and ABX using a total of 99 subjects, control, CRSsNP, and CRSwNP patients, respectively (horizontal line = median and whiskers = min/max range). (F) LEfSe analysis identified NABX-enriched families (colored in blue) and ABX-enriched families (colored in red) in CRSwNP patients. The plot indicates taxa with an LDA score > 3.0 and a *p* < 0.05 in all-against-all comparisons (more stringent). (\**p* < 0.05, \*\**p* < 0.01, \*\*\**p* < 0.001)

Next, we compared the nasal bacterial composition in NABX and ABX. In NABX, *Firmicute, Cyanobacteria*, and *Bacteroidetes* significantly differed according to disease status (Fig. S5A and C). At the family level, the relative abundance of *Propionibacteriaceae, Moraxellaceae*, and *Prevotellaceae* significantly differed according to disease status in NABX (Fig. 2C and Fig. S6A). LEfSe analysis revealed that *Propionibacteriaceae* was significantly decreased in CRSwNP and CRSsNP compared to levels in the control group (Fig. S6B). In the ABX, there was no significant difference in phyla and families that possessed a relative abundance of greater than 3 percent (Fig. 2D and Fig. S5B). When all identified families were analyzed using LEfSe, we observed that *Peptostreptococcaceae, Desulfovibrionaceae*, and *Alteromonadaceae* were the most dominant families in the control group (Fig. S6C). *Sulfurovaceae* was found to be significantly enriched in CRSsNP. Taken together, these findings indicate that antibiotic use could reduce differences in microbial communities according to disease status.

To determine the differences in the microbial community resulting from antibiotic use, we compared NABX and ABX in regard to each disease status. In a total of 99 subjects, unlike alpha diversities, PERMANOVA revealed that microbial composition significantly differed between NABX and ABX (Fig. 2E and Fig. S7A and B). Among bacterial families with a relative abundance of more than 3%, *Staphylococcaceae, Propionibacteriaceae*, and *Streptococcaceae* exhibited significant differences between the two groups (Fig. S7C). In all families that we identified, we found that *Staphylococcaceae, Streptococcaceae*, and *Propionibacteriaceae* were significantly decreased in ABX compared to levels in NABX (Fig. S7D). *Entotheonellaceae* and *Sandaracinaceae* were significantly enriched in ABX compared to levels in NABX.

Then, we compared the microbial communities between ABX and NABX in control group. There was no significant difference in alpha and beta diversities between the two groups (Fig. 2E and Fig. S8A and B). Additionally, families with a relative abundance of more than 3% were not significantly different between the two groups (Fig. S8C). LEfSe analysis indicated that antibiotic use led to enrichment of families whose relative abundance was lower than 3% (Fig. S8D). In CRSsNP patients, the alpha diversity indices and microbial composition did not significantly differ between NABX and ABX (Fig. 2E and Fig. S9A and B). Among the families with a relative abundance of more than 3%, *Staphylococcaceae* was significantly decreased in ABX compared to levels in NABX (Fig. S9C). LEfSe analysis that was performed on all identified families revealed that *Staphylococcaceae, Intrasporangiaceae*, and *Neisseriaceae* were significantly decreased in ABX compared to levels in NABX (Fig. S9D). Additionally, *Prevotellaceae* and *Legionellaceae* were enriched significantly in ABX compared to levels in NABX. Last, we compared the microbial communities between NABX and ABX in CRSwNP patients. Unlike Chao1 and the number of observed OTUs, Shannon and Simpson indices were significantly lower in ABX compared to those in NABX (Fig. 2E and Fig. S10A). Additionally, PERMANOVA revealed significant differences in microbial composition between the two groups (Fig. S10B). According to these results, it appeared that antibiotics exerted stronger effects on the microbial community in CRSwNP patients (NP) compared to those in the control and CRSsNP patients (Non-NP). Among the bacterial families with more than 3% relative abundance, *Streptococcaceae* and *Lachnospiraceae* were significantly decreased in the ABX compared to levels in the NABX group (Fig. S10C). In all identified families, LEfSe analysis revealed that *Streptococcaceae, Lachnospiraceae*, and *Neisseriaceae* were significantly decreased in ABX compared to levels in NABX (Fig. 2F).

### Differences in secreted proteome according to the use of antibiotics

As we determined, antibiotics may exert stronger effects on the microbial community in NP compared to that in Non-NP groups, and based on this, we also compared host responses to antibiotics in the NP and Non-NP groups using proteomic analysis. The NABX group consisted of 22 control, 19 CRSsNP, and 11 CRSwNP patients, while the ABX group consisted of 1 control, 5 CRSsNP, and 11 CRSwNP patients. In the ABX group, relatively small proteomic changes were observed between NP and Non-NP patients compared to those in the NABX group (Fig. 3A). This was consistent with our results indicating that the use of antibiotics reduced differences in microbial communities according to disease status. As shown in Tables S2 and S3, proteins with a fold change ≥ 2.0 at a *p* value < 0.05 in the NABX group were considerably different compared to those in the ABX group. Furthermore, to determine the differences in the secreted proteome based on antibiotic use, we compared NABX and ABX in 69 patients, Non-NP and NP (Fig. 3B and Table S4-6). Based on previous results obtained using metagenomics, antibiotics may exert stronger effects on the secreted proteome in NP patients compared to that in total and Non-NP patients. To identify the canonical pathways that were most significantly involved with the proteins that exhibited a fold change ≥ 2.0 at a *p* value < 0.05 in each group, we performed ingenuity pathway analysis (IPA). When examining all subjects, the pathways were completely different from those identified in NP, as the NP pathways were associated with innate immunity and production NO and ROS (Fig. 3C and D). There was no significant pathway found in the Non-NP group (*p* > 0.001). As LXL/RXR activation was the most significant pathway according to IPA analysis, proteins associated this pathway were labeled in a volcano plot derived from the NP group (Fig. 3B). These analyses indicated that the use of antibiotics may result in stronger effects on the nasal microbiome and the secreted proteome in NP compared to those in Non-NP.

**Figure 3.**
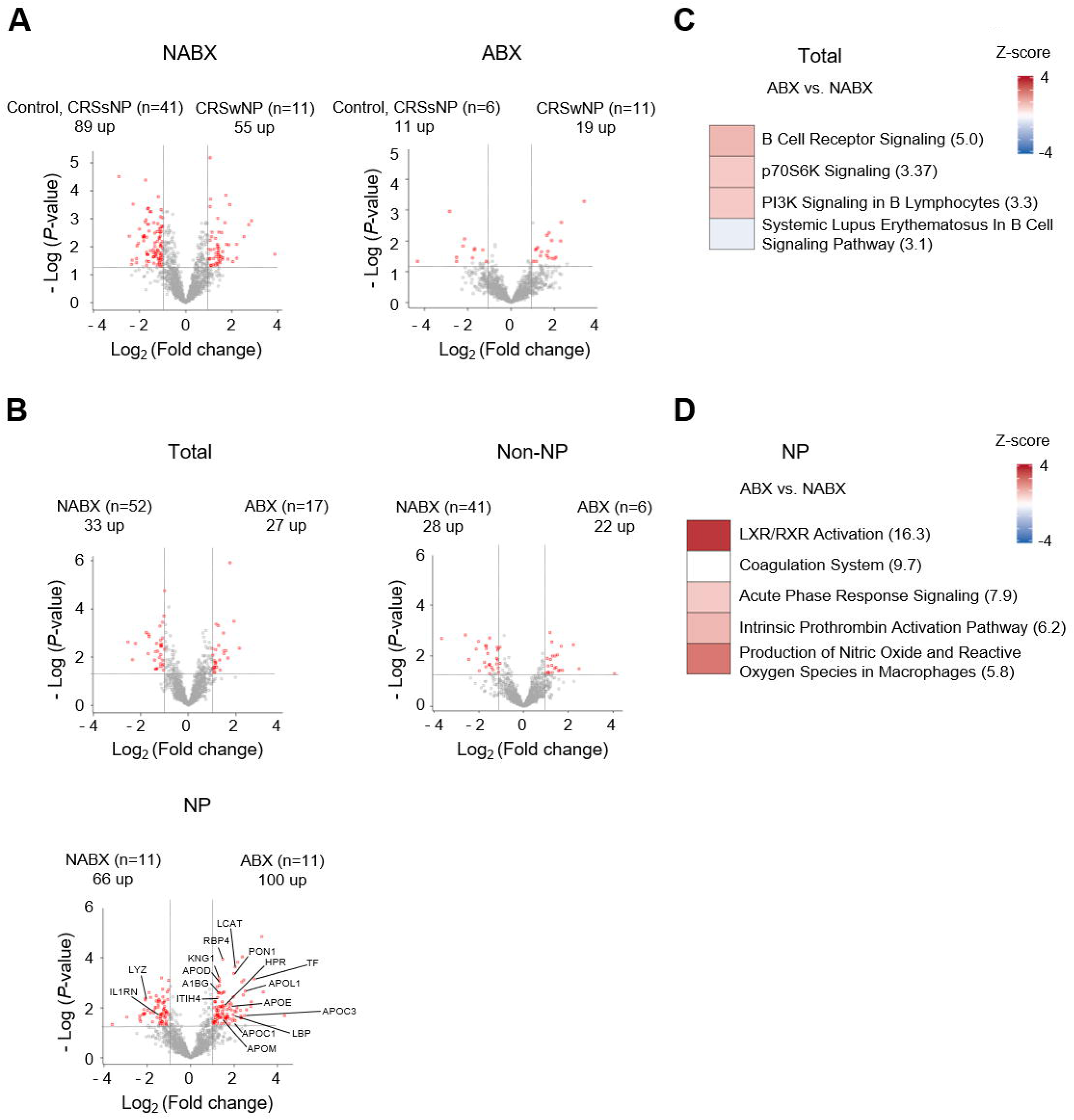
Differences in the secreted proteome according to the use of antibiotics. (A and B) Volcano plots of the secreted proteome where log_2_ of fold changes was plotted versus the - log_10_ of the *p*-values from the Student’s t-test. The horizontal dashed line indicates *p*-values of 0.05, and the vertical dashed lines indicate fold changes of 2.0. Red squares represent proteins with a fold change ≥ 2.0 at *p* value < 0.05. In NP, the labeled squares indicate proteins involved in LXR/RXR activation. (C and D) Canonical pathway analysis representing significantly up- or down-regulated pathways in ABX compared to those in NABX. The pathways with *p*-values smaller than 0.001 were arranged from top to bottom in order of the lowest to the highest *p*-value. The parenthesis represented the -log_10_ *p*-value.

### Antibiotic-dependent relationships between nasal microbiome and secreted proteome

We next sought to investigate if the associations between the microbiome and the secreted proteome may differ according to antibiotic use. If associations with a high number of significant correlations were altered by exposure to antibiotics, this would be meaningful. Therefore, we arranged the microbiome and secreted proteome in descending order from the highest to the lowest number of significant correlations with each other (Fig. S11A and B). Among these, we clustered the top 25 percent of the microbiome and secreted proteome in NABX and ABX (Fig. 4A and B). The average *R*-squared value in ABX was higher than that in NABX. From these analyses, we confirmed that the microbiome and secreted proteome in ABX correlated strongly with one another compared to the correlation observed in NABX. Furthermore, among these top 25 percent microbiome and secreted proteome components in NABX and ABX, there was little overlap between NABX and ABX (Fig. 4C and D). Likewise, the use of antibiotics altered the associations between the microbiome and secreted proteome. Moreover, their correlations were strengthened in subjects who had taken antibiotics.

**Figure 4.**
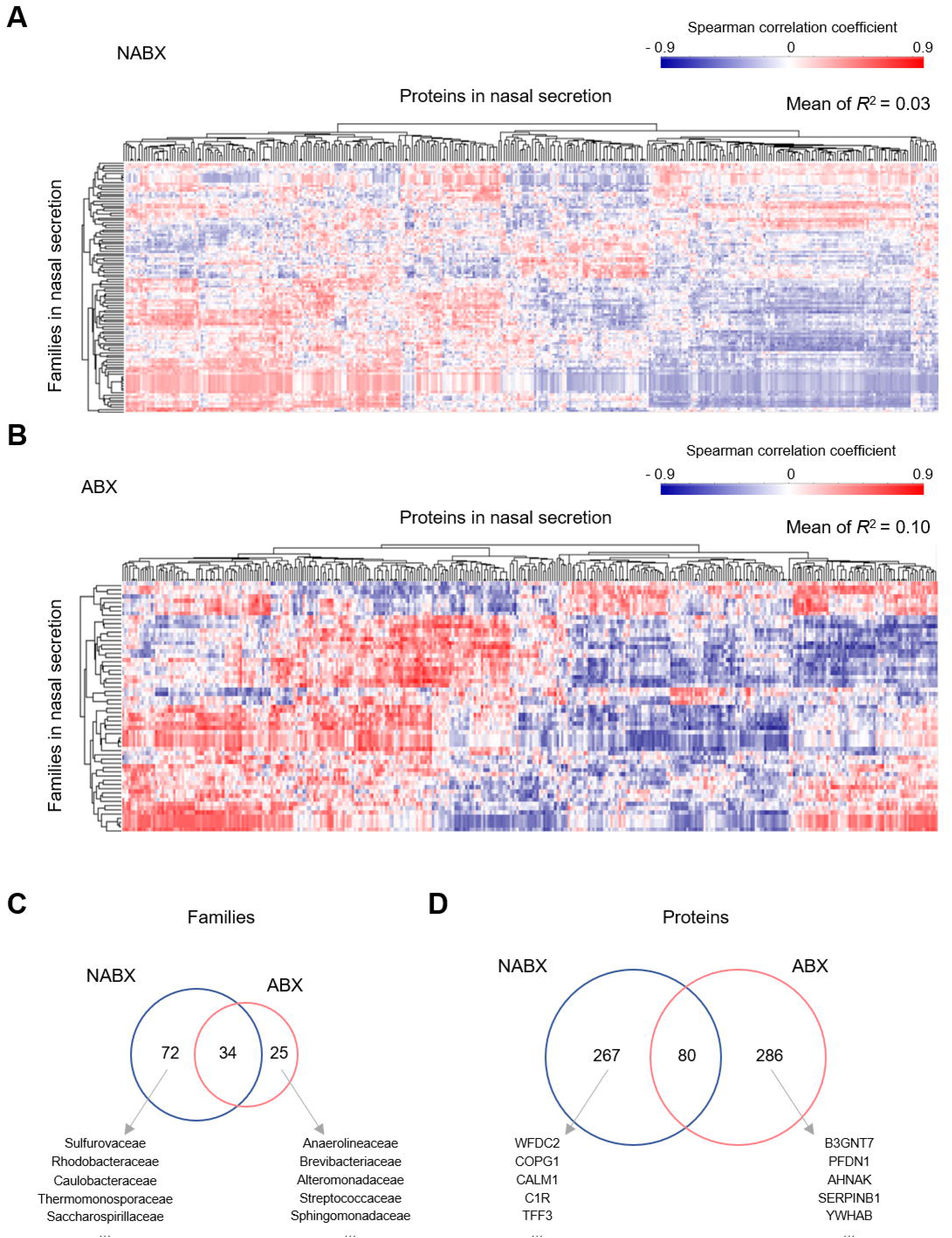
Antibiotic-dependent relationships between the nasal microbiome and the secreted proteome. (A and B) Hierarchical clustering of the top 25 percent of the microbiome and proteome with a high number of significant correlations in NABX and ABX. The means of squared coefficients were calculated using all values presented in (A and B). Among the microbiomes and proteomes identified in (A and B), a Venn diagram reveals the number of families (C) and proteins (D) in NABX and ABX. The top 5 proteins and families (non-overlapping and non-redundant) are represented.

### Antibiotic-dependent relationships between the nasal microbiome and the secreted proteome in Non-NP and NP

Finally, given that antibiotics may exert different effects on the nasal microbiome, secreted proteome, and their association according to disease status, we hypothesized that the associations that were altered by antibiotics could differ according to disease status. To confirm this hypothesis, we divided the nasal microbiome into two groups that included families with increased relative abundance in ABX (IFa) and families with decreased relative abundance in ABX (DFa). We also divided the secreted proteome into two groups that included proteins with increased normalized intensity in ABX (IPa) and proteins with decreased normalized intensity in ABX (DPa). We could not detect the IFa group in the NP patients. A number of significant positive or negative correlations were observed among the four groups (IFa, DFa, IPa and DPa) in both the Non-NP and NP patient groups (Fig. 5A and B). We confirmed that NP patients showed stronger correlations with each other than those in the Non-NP patients. To confirm the associations among the four groups, we performed aSPC tests (Fig. 5C). The associations between DFa and IPa or DFa and DPa in the NP patients were more significant than those observed in the Non-NP patients; however, we found no correlation between each group (IFa, DFa, IPa and DPa) and others in regard to the microbiome and the proteome. From these analyses, in NP it was apparent that the associations between the microbiome and the proteome that were altered by antibiotics were different compared to those in the Non-NP patients. Moreover, the associations among the four groups in the NP patients were more significant than those in the Non-NP patients.

**Figure 5.**
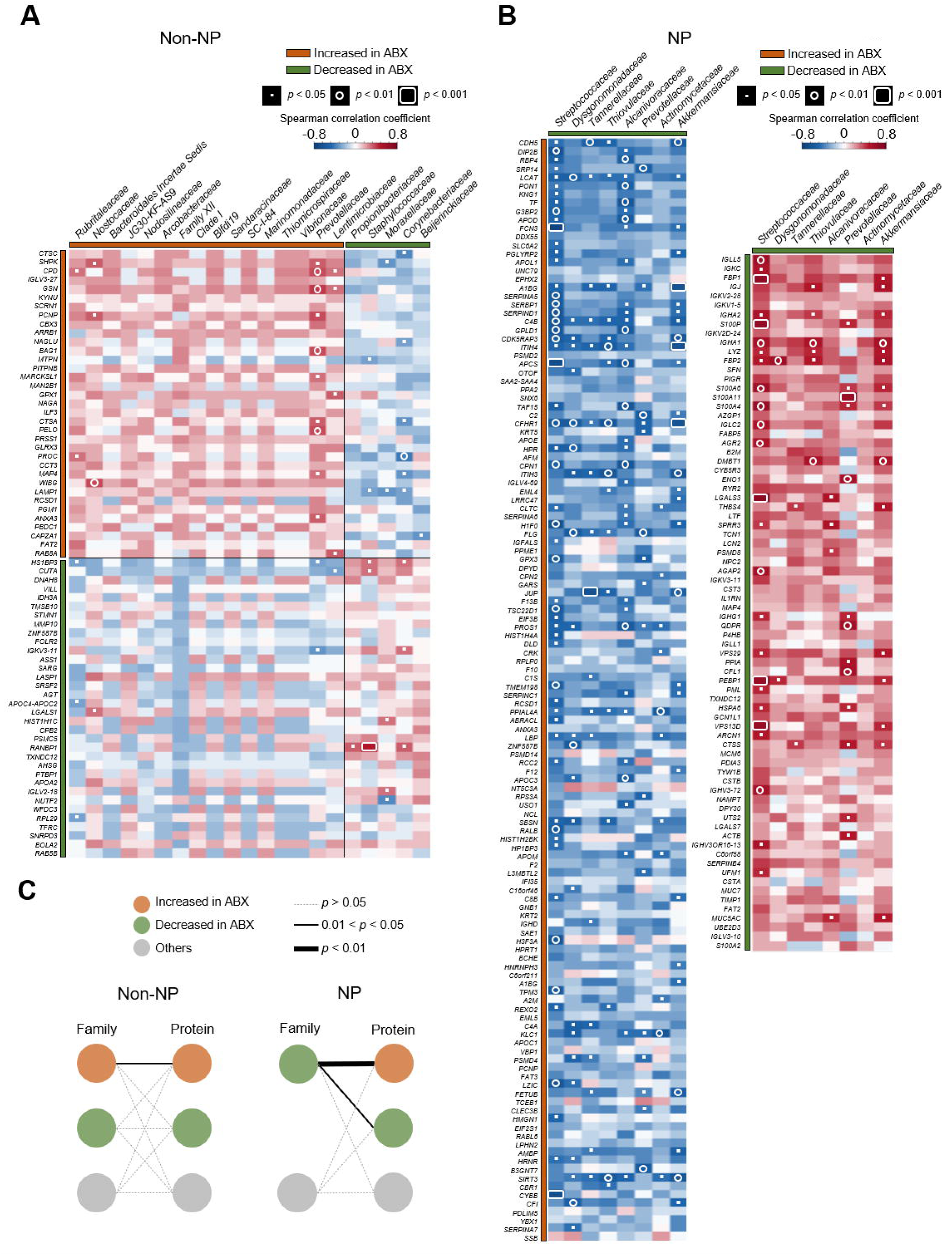
Antibiotic-dependent relationships between the nasal microbiome and the secreted proteome in control, CRSsNP, and CRSwNP patients. (A and B) Spearman correlation heatmaps of the secreted proteomes (rows) and the nasal microbiomes (columns) that exhibit significant differences between NABX and ABX in Non-NP and NP groups. The proteome (Student’s t-test, *p* < 0.05) and microbiome (Mann-Whitney U test, *p* < 0.05) were arranged from top to bottom and from left to right in order of the lowest to the highest *p*-value. Orange and green boxes indicate families and proteins enriched in ABX and NABX. Redundant microbiomes and proteomes were excluded from the heatmap. (C) Associations between the microbiome and the proteome. Orange and green circles represent the same groups as described in (A and B), respectively. Gray circles indicate other microbiomes and proteomes that were identified in this study (adaptive sum of powered correlation [aSPC] test, *p* < 0.05).

## Discussion

To our knowledge, this is the first report to analyze the effects of antibiotics on the microbiome, secreted proteome, and their associations in CRS using multi-omics. The relatively large number of subjects used for this study enabled us to more fully understand the associations between CRS and antibiotics. As expected, antibiotic use could reduce differences in the microbial community and the secreted proteome according to disease status. Interestingly, antibiotics may exert strong effects on not only the nasal microbiome and secreted proteome, but also their associations in NP patients compared to that in Non-NP patients. Additionally, their correlations were strengthened in subjects who had taken antibiotics.

Previous studies examining the nasal microbiome resulted in inconsistent outcomes for alpha diversity indices in CRS patients compared to those in the control group. Some studies reported that there was no significant difference in alpha diversity between the control and the CRS patients(Koeller et al. 2018; Mahdavinia et al. 2018; Paramasivan et al. 2019). Meanwhile, other studies reported that the alpha diversity indices significantly decreased in CRS patients compared to those in the control group(Choi et al. 2014; Biswas et al. 2015; Cope et al. 2017). However, Aurora *et al*.(Aurora et al. 2013) reported that bacterial diversity was increased in CRS compared to that in the control. Additionally, Lal *et al*.(Lal et al. 2017) identified that control and CRSwNP patients exhibited significantly higher faith phylogenic and Shannon diversity compared to those values in CRSsNP patients. Copeland *et al*.(Copeland et al. 2018) reported that Shannon diversity was significantly increased in CRSsNP patients compared to that in control and CRSwNP patients. Moreover, a number of previous studies(Biswas et al. 2017; Hoggard et al. 2017; Chalermwatanachai et al. 2018; Gan et al. 2019; Rom et al. 2019) indicated that alpha diversity was increased, decreased, or unchanged in CRS compared to that in control, and this was dependent upon the alpha diversity index.

To date, the existence of any association between antibiotic use and nasal microbiome in the context of CRS remains unknown. Palleja *et al*.(Palleja et al. 2018) demonstrated that microbial richness within the gut was significantly decreased at approximately 6 months following antibiotic treatment. As the nasal microbiome studies in CRS generally excluded patients who had taken antibiotics within approximately 1 month prior to sampling, the differences in alpha diversity according disease status could not be clearly identified. In our study that included subjects who had taken antibiotics within three months, Shannon and Simpson indices were significantly increased from control to CRSwNP patients, and we are confident that these results reflect real-world criteria. Based on this, antibiotic use should be taken into account when comparing diversity according to disease status.

We also found that antibiotics could exert stronger effects in NP compared to that in Non-NP patients. Additionally, antibiotics could alter the nasal secreted proteome and the associations between the microbiome and the secreted proteome. However, the mechanisms by which antibiotics could affect the microbiome according to disease status and could alter the secreted proteome remain unknown. Shaw *et al*.(Shaw et al. 2019) reported a simple quantitative model based on a stability landscape framework. They demonstrated that the microbiome existed in multiple stable equilibria of landscape that could be influenced by sufficiently strong perturbations such as antibiotics that could alter the microbiome from its normal equilibrium to other states. We speculated that perturbation by antibiotics could shift the microbiome towards equilibrium possessing a different value of alpha diversity (lower diversity) in only NP patients. On the other hands, treatment with antibiotics disrupted the intestinal tight junction(Tulstrup et al. 2015; Feng et al. 2019). Kohanski *et al*.(Kohanski et al. 2017) reported that levofloxacin, a commonly used antibiotic for upper airway infections, simulated reactive oxygen species (ROS) and caspase-3 activity in cultured human sinonasal epithelial cells. Thus, we speculated that antibiotic use could affect the nasal epithelial cells and alter the proteome secreted by these cells.

The IPA analysis revealed that B cell-associated pathways and anti-microbial pathways were more highly activated in ABX patients compared to levels in NABX patients in the total patient population and the NP group, respectively. PI3K regulated B cell receptor-mediated antigen presentation and airway remodeling(Yoo et al. 2017). Activated B cells utilized mTORC1 to activate p70 ribosomal S6 kinase (p70S6K), and this preceded antibody synthesis(Gaudette et al. 2020). In contrast, macrophages that released proinflammatory cytokines played an important role in clearing bacteria, and this was accompanied by the production of nitric oxide (NO) and ROS upon contact with pathogens(Tan et al. 2016; Finney et al. 2019). Several studies have reported that LXR activation exerted anti-inflammatory functions(Higham et al. 2013) and inhibited bacterial infection of host macrophages(Pascual-Garcia et al. 2013; Matalonga et al. 2017).

There were limitations in this present research study. First, the proportion of NABX and ABX in the control group was smaller than that in the CRSsNP and CRSwNP groups. Second, larger number of subjects should be analyzed to improve reliability. Thus, a similar proportion of subjects who had taken antibiotics and a larger sample size for each disease status will be required to verify the association between CRS and antibiotics.

In summary, we determined that antibiotics could exert strong effects on the microbial diversity and the secreted proteome in NP patients compared to those in Non-NP patients. Integrative analyses revealed that the associations and correlations between the microbiome and the secreted proteome could be altered and strengthened in subjects who had recently used antibiotics. These findings provide new insight into the effects of antibiotics on the nasal environment and the host response in CRS.

## Methods

### Study design and collection of nasal secretions

The Internal Review Board of Seoul National University Hospital (SNUH) (no. C-1308-099-515) approved this study. All subjects were informed of the purpose of the study, and they all signed written informed consent forms. The diagnosis of CRS was based on patient history, physical examination, nasal endoscopy, and sinus computed tomography in accordance with the 2012 European position paper on rhinosinusitis and nasal polyps (EPOS) guidelines(Fokkens et al. 2012). Patients possessing deviated nasal septa but lacking any sinonasal disease were considered as the control. Subjects that were less than 14 years of age and who were diagnosed with unilateral rhinosinusitis, antrochoanal polyps, allergic fungal sinusitis, cystic fibrosis, or immotile ciliary disease were excluded from the study. The demographic characteristics of the subjects are summarized in Table 1. Nasal secretions were obtained from both sides of the nose as previously described(Kim et al. 2019). Sterilized strips of filter paper (7 × 30 mm; Whatman No. 42, Whatman, Clifton, NJ, USA) were placed on the middle meatus for 10 minutes. Two filter papers from each subject were transferred into a tube. Then, 1 mL of nuclease-free water was added to each tube, and the tubes were rotated for 1 h at room temperature. The nasal secretions were stored in aliquots at -70 °C. The workflow is shown in Figure S1.

**Table 1.**
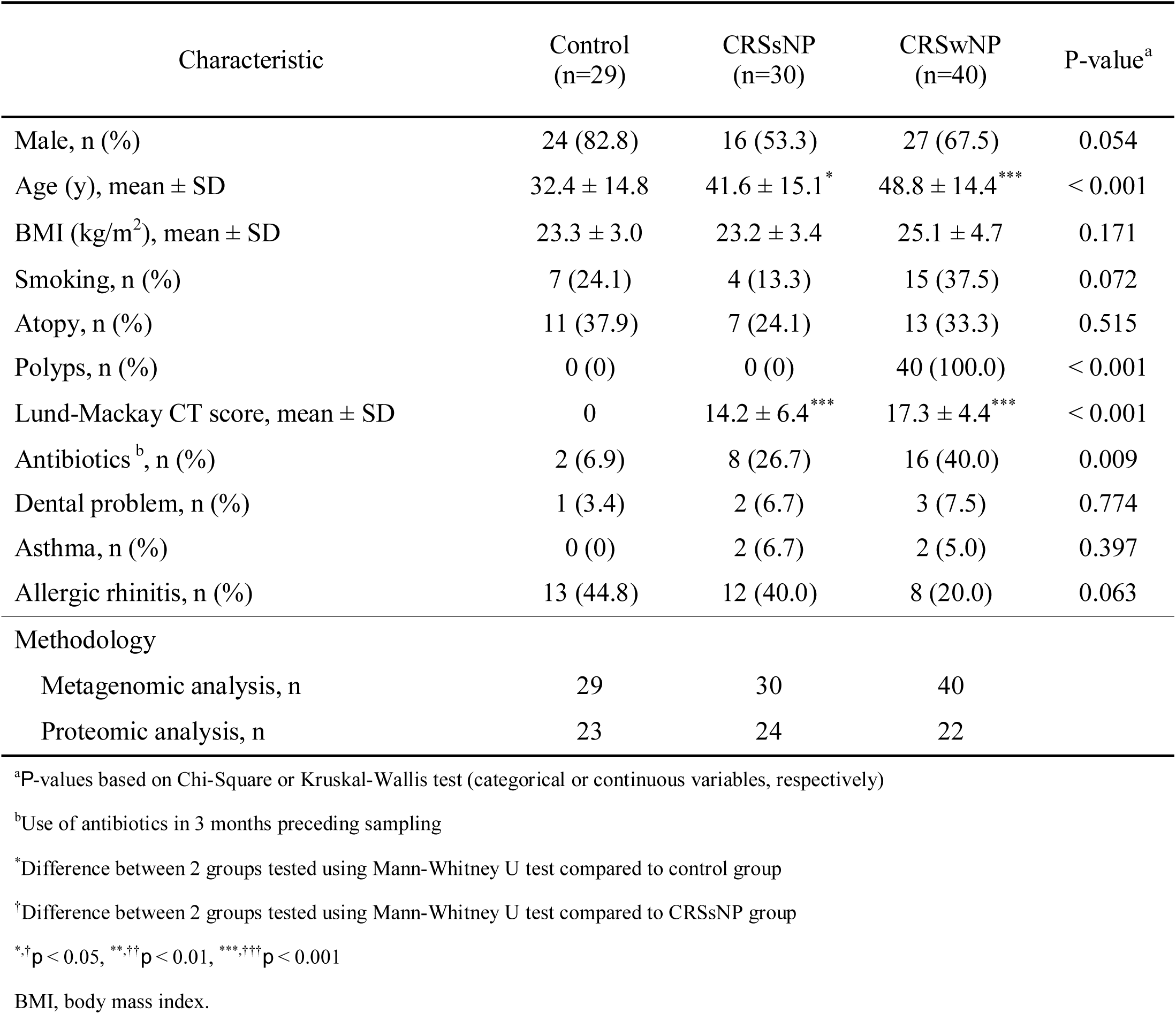
Demographic characteristics of subjects.

### DNA extraction and sequencing

After preparation of nasal secretions, 16S rDNA was extracted using the PowerSoil DNA Isolation Kit (Mo Bio Laboratories, Inc., Carlsbad, CA, USA). The manufacturer’s instructions for the kit were followed for all subsequent procedures. The isolated DNA was sent to Macrogen Corporation (Macrogen Inc., Seoul, South Korea) for amplification and sequenced on a Miseq instrument (Illumina, USA). A sequencing library was prepared by amplifying the V3-4 region of 16S rDNA. For the amplification, primers 341F (5’ -TCG TCG GCA GCG TCA GAT GTG TAT AAG AGA CAG CCT ACG GGN GGC WGC A-3’) and 805R (5’ -GTC TCG TGG GCT CGG AGA TGT GTA TAA GAG ACA GGA CTA CHV GGG TAT CTA ATC C-3’) were used. The library was quantified using TapeStation DNA ScreenTape D1000 (Agilent, USA) and Picogreen assay. The 16S rDNA libraries were sequenced using the MiSeq platform for 2 × 300 cycles.

### Preparation of nasal secretions for proteomics

A 100 μL aliquot of each nasal secretion was prepared as previously described(Kim et al. 2019). The samples were centrifuged at 15,000 rpm for 10 minutes at 4 °C. We measured the protein concentration in the supernatant by tryptophan fluorescence emission at 350 nm using an excitation wavelength of 295 nm(Wisniewski and Gaugaz 2015). For the analysis, 50 μg of nasal secreted proteins were used per sample. Proteins were digested using the 2-step FASP procedure with some modifications(Han et al. 2014; Woo et al. 2015). Pellets were resuspended in SDT buffer (2% SDS, 10mM TCEP, and 50mM CAA in 0.1M Tris pH 8.0). Then, the solution loaded onto a 10K Amicon filter (Milipore). The buffer was exchanged to UA solution (8M urea in 0.1M Tris pH 8.5) using centrifugation at 14,000 g. Following the exchange of 40mM ammonium bicarbonate (ABC) buffer, proteins were digested at 37 °C overnight using a trypsin/LysC mixture (protein-to-protease ratio of 100:1). The peptides were isolated using centrifugation. After the filter units were rinsed with 40 mM ABC, we performed second digestion at 37 °C for 2 hours using trypsin (enzyme-to-substrate ratio [w/w] of 1:1000). The digested peptides were acidified using 10% TFA and desalted using homemade C18-StageTips as described(Han et al. 2014; Woo et al. 2015). Finally, we used a vacuum dryer to dry it and stored at -80 °C.

### High-pH StageTip-based peptide fractionation

For peptide spectrum library, we performed StageTip-based, high-pH peptide fractionation as described with some modifications(Han et al. 2014). Peptides obtained from pooled samples were dissolved in 200 μL of loading solution (10 mM ammonium hydroxide solution, pH 10 and 2% acetonitrile) and separated on the reversed-phase tip columns, prepared by packing POROS 20 R2 (Invitrogen, Carlsbad, CA) into a 200 μL yellow tip with C18 Empore disk membranes (3M, Bracknell, UK) at the bottom. After conditioning of microcolumns with methanol, acetonitrile, and loading buffer, peptides were loaded at pH 10, and 20 fractions were subsequently eluted with buffer solutions, pH 10, containing 5%, 10% 15%, 20%, 25%, 30%, 35%, 40%, 60%, and 80% acetonitrile. To improve the orthogonal fractionation of the RP-RP separation, 20 fractions were combined into six fractions in a noncontiguous manner. The six fractions were dried in a vacuum centrifuge and stored at -80 °C until LC-MS/MS analysis.

### LC-MS/MS analysis

Liquid chromatography (LC)-mass spectrometry (MS)/MS analysis was performed using a Q-exactive plus (Thermo Fisher Scientific, Waltham, MA) coupled to an Ultimate 3000 RSLC system (Dionex) via a nano electrospray source as described with some modifications(Lee et al. 2018; Kim et al. 2019). Peptides in samples were separated using the 2-column setup (a trap column [300 μm I.D. × 5 mm, C18 3 μm, 100 Å] and an analytical column [50 μm I.D. × 50 cm, C18 1.9 μm, 100 Å]). Prior to injection of each sample, we resuspended the dried peptides in solvent A (2% acetonitrile and 0.1% formic acid). After the samples were loaded onto the nano LC, the samples were run with a 90 minute gradient from 8% to 30% solvent B (80% acetonitrile and 0.1% formic acid). The spray voltage was set to 2.0 kV in the positive ion mode, and the temperature of the heated capillary was 320 °C. The MS method consisted of a survey scan at 35,000 resolution from 400 to 1,220 m/z. Automatic gain control (AGC) target of 3×10^6^ at injection time of 60ms. Then, 19 DIA windows were acquired at 35,000 resolution with AGC target 3×10^6^ and auto for injection time(Bruderer et al. 2015). Stepped collision energy (SCE) was 10% at 27%.

### Data processing for spectral library construction

First, we processed MS raw files that obtained from 18 data-dependent acquisition (DDA) measurements of the pooled samples using MaxQuant (version 1.6.1.0)(Tyanova et al. 2016a). MS/MS spectra were searched against the Human Uniprot database (December 2014, 88,657 entries) and the Biognosys iRT peptides fasta database using the Andromeda(Cox et al. 2011). Data was searched with 6 ppm precursor ion tolerance for total protein level analysis and 20 ppm MS/MS ion tolerance. We used variable modifications (N-acetylation of protein and oxidation of methionine) and a fixed modification (cysteine carbamido-methylation). Enzyme specificity was set as full tryptic digestion. Peptides with a minimum length of six amino-acids and up to two missed-cleavages were considered. The required false discovery rate (FDR) was set to 1% at the peptide, protein, and modification level. To maximize the number of quantification events across samples, we enabled the ‘Match between Runs’ option on the MaxQuant platform. MaxQuant search results were imported as spectral libraries into Spectronaut Pulsar X with default settings.

In addition to the typical DDA-based spectral library, we generated a spectral library from the DIA data using the Spectronaut software (Biognosys, Schlieren, Switzerland). The Pulsar search engine integrated in Spectronaut Pulsar X was used to identify peptides and proteins using only independent DIA dataset with the same search engine parameters (fasta database, modifications, peptide length, miss-cleavage, peptide, and protein FDR) as listed above for MaxQuant.

### Data processing for the DIA MS

The DIA data of individual samples were analyzed with Spectronaut Pulsar X (Biognosys, Schlieren, Switzerland). First, we converted the DIA raw files into an htrm format by using the GTRMS Converter provided by the Spectranaut. The FDR was estimated with the mProphet(Reiter et al. 2011) approach and set to 1% at peptide precursor level and at 1% at protein level. The proteins were inferred by the software, and the quantification information was acquired at the protein level by using the q-value < 0.01 criteria, which was used for subsequent analyses.

### Bioinformatic analysis

Raw sequences were analyzed and quality-filtered using Quantitative Insights Into Microbial Ecology (QIIME) (version 1.9.1)(Caporaso et al. 2010). An operational taxonomic unit (OTU) was defined at 97% sequence identity using UCLUST against the SILVA reference sequence database (version 132)(Edgar 2010; Quast et al. 2013). Singletons were removed from the OTU table to reduce the noise and finally, 12,634,938 sequences remained. Alpha diversity (Chao1, the number of observed OTUs, Shannon, and Simpson) was calculated in QIIME. Beta diversity was calculated using a Bray–Curtis distance matrix with the vegan package in R software. In addition, principal coordinates analysis (PCoA) was performed using the Phyloseq package in R software(McMurdie and Holmes 2013). Linear discriminant analysis (LDA) effect size (LEfSe) was performed to determine bacterial features most likely to describe differences between groups among all of the identified bacterial families(Segata et al. 2011). The relative abundance of each OTU was calculated by dividing the read counts of an OTU by the total read counts of all OTUs in each subject, except for read counts of *Archaea, Chloroplast* and *Mitochondria*. Proteomic data were normalized using width adjustment. Ingenuity Pathway Analysis (IPA; QUAGEN, Hilden, Germany) software was used to predict canonical pathways. Missing values in proteomic analysis were imputed by normal distribution (width = 0.15, downshift = 1.8) using Perseus(Tyanova et al. 2016b). Graphical outputs were visualized using R software (version 3.5.1), Perseus (version 1.6.0), and GraphPad Prism (version 8.1.1).

### Statistical analysis

Student’s t-test or ANOVA was performed to compare the data under normal distribution, while Mann-Whitney *U* or Kruskal-Wallis tests were used for non-parametric analysis. A Chi-square test was performed to examine the associations between categorical variables. The significant differences between groups were calculated using permutational multivariate analysis of variance (PERMANOVA) via Adonis function in vegan package in R software. LEfSe analysis used the Wilcoxon rank-sum test or Kruskal-Wallis rank-sum test. In IPA analysis, *p*-value of Fisher’s exact test < 0.001 was considered significant pathways. Spearman correlation was used to assess the correlation and adaptive sum of powered correlation test (aSPC)(Xu et al. 2017) was performed to calculate global association between microbiome and proteome. The statistical analyses were conducted using SPSS ver. 25.0 (SPSS Inc., Chicago, IL, USA), Perseus (version 1.6.0), and R software (version 3.5.1).

## Abbreviations

CRS: Chronic rhinosinusitis;
CRSsNP: Chronic rhinosinusitis without nasal polyp;
CRSwNP: Chronic rhinosinusitis with nasal polyp;
ABX: the subjects who had taken antibiotics 3 months before sampling;
NABX: the subjects who had not taken antibiotics 3 months before sampling;
L-M: Lund-Mackay;
SPT: skin prick test;
PERMANOVA: Permutational multivariate analysis of variance;
PCoA: principal coordinates analysis;
LDA: Linear discriminant analysis;
LEfSe: Linear discriminant analysis effect size;
OTU: Operational taxonomic unit;
LC-MS: Liquid chromatography-mass spectrometry;
DIA: Data-independent acquisition;
IPA: Ingenuity pathway analysis;
NP: Nasal polyp;
aSPC: adaptive sum of powered correlation

## Author’s contributions

Conception and design: Yi-Sook Kim, Hyun-Woo Shin; Acquisition of data: Yi-Sook Kim, Dohyun Han, Ji-Hun Mo, Yong-Min Kim, Dae Woo Kim, Hyo-Guen Choi; Analysis and interpretation of data: Yi-Sook Kim, Dohyun Han, Hyun-Woo Shin; Drafting the article or revising it critically for important intellectual content: Yi-Sook Kim, Jong-Wan Park, Hyun-Woo Shin; Supervision: Hyun-Woo Shin

## Availability of data and materials

The raw sequence data for the metagenomics are available in the NCBI Sequence Read Archive at study accession number PRJNA557492.

