## Supplemental figure legends and tables for "Antibiotic-dependent relationships between nasal microbiome and secreted proteome in chronic rhinosinusitis and nasal polyps"

**Supplementary figures**

**Figure S1. Overview of the study**

**Figure S2. Comparison of diversity and microbial composition among control, CRSsNP, and CRSwNP**

(A) Comparison of Chao1 and the number of observed OTUs between disease status (horizontal line = median and whiskers = min/max range). (B) Distribution of bacterial phyla between disease status. The composition of each phylum with relative abundance of more than 3 percent were illustrated. Among the taxa with relative abundance of more than 3 percent, the dot plots showed relative abundance of phyla (C) and families (D) with significant differences between disease status (horizontal line = mean). (E) LEfSe analysis identified control-enriched families (colored in blue), CRSsNP-enriched families (colored in green), and CRSwNP-enriched families (colored in red). The plot showed taxa with LDA score > 3.0 and *p* < 0.05 in all-against-all (more stringent). (Kruskal-Wallis and Mann-Whitney U test, **p* < 0.05, ***p* < 0.01, ****p* < 0.001)

**Figure S3. Number of the quantified proteins in each group**

A bar plot of total number of the quantified proteins from the two technical replicates in proteomics. Error bars represented the standard deviation of the mean.

**Figure S4. Comparison of microbial composition between disease status according to the use of antibiotics**

(A and B) Comparison of Chao1 and the number of observed OTUs between disease status in NABX and ABX (horizontal line = median and whiskers = min/max range). (C and D) PCoA plots based on Bray–Curtis distance matrix in NABX and ABX.

**Figure S5. Phylum composition between disease status according to the use of antibiotics**

(A and B) Distribution of bacterial phyla between disease status. The composition of each phylum with relative abundance of more than 3 percent were illustrated in NABX and ABX. (C) Among the taxa with relative abundance of more than 3 percent, the dot plot showed relative abundance of phyla with significant differences between disease status in NABX (horizontal line = mean). (**p* < 0.05, ***p* < 0.01, ****p* < 0.001)

**Figure S6. Family composition between disease status according to the use of antibiotics**

(A) Among the taxa with relative abundance of more than 3 percent, the dot plot showed relative abundance of families with significant differences between disease status in NABX (horizontal line = mean). (B and C) LEfSe analysis identified control-enriched families (colored in blue), CRSsNP-enriched families (colored in green), and CRSwNP-enriched families (colored in red) in NABX and ABX. The plot showed taxa with LDA score > 3.0 and *p* < 0.05 in all-against-all (more stringent). (**p* < 0.05, ***p* < 0.01, ****p* < 0.001)

**Figure S7. Differences in microbial composition according to the use of antibiotics** **in total of 99 subjects**

(A) Comparison of Chao1 and the number of observed OTUs between NABX and ABX in 99 patients (horizontal line = median and whiskers = min/max range). (B) A PCoA plot based on Bray–Curtis distance matrix. (C) Distribution of bacterial families between NABX and ABX. The composition of each family with relative abundance of more than 3 percent were illustrated. The parenthesis indicated phylum belonged to the family. (D) LEfSe analysis identified NABX-enriched families (colored in blue) and ABX-enriched families (colored in red). The plot showed taxa with LDA score > 3.0 and *p* < 0.05 in all-against-all (more stringent). (**p* < 0.05, ***p* < 0.01, ****p* < 0.001)

**Figure S8. Differences in microbial composition according to the use of antibiotics** **in control**

(A) Comparison of Chao1 and the number of observed OTUs between NABX and ABX in control (horizontal line = median and whiskers = min/max range). (B) A PCoA plot based on Bray–Curtis distance matrix. (C) Distribution of bacterial families between NABX and ABX. The composition of each family with relative abundance of more than 3 percent were illustrated. The parenthesis indicated phylum belonged to the family. (D) LEfSe analysis identified NABX-enriched families (colored in blue) and ABX-enriched families (colored in red). The plot showed taxa with LDA score > 3.0 and *p* < 0.05 in all-against-all (more stringent).

**Figure S9. Differences in microbial composition according to the use of antibiotics** **in CRSsNP**

(A) Comparison of Chao1 and the number of observed OTUs between NABX and ABX in CRSsNP (horizontal line = median and whiskers = min/max range). (B) A PCoA plot based on Bray–Curtis distance matrix. (C) Distribution of bacterial families between NABX and ABX. The composition of each family with relative abundance of more than 3 percent were illustrated. The parenthesis indicated phylum belonged to the family. (D) LEfSe analysis identified NABX-enriched families (colored in blue) and ABX-enriched families (colored in red). The plot showed taxa with LDA score > 3.0 and *p* < 0.05 in all-against-all (more stringent). (**p* < 0.05, ***p* < 0.01, ****p* < 0.001)

**Figure S10. Differences in microbial composition according to the use of antibiotics** **in CRSwNP**

(A) Comparison of Chao1 and the number of observed OTUs between NABX and ABX in CRSwNP (horizontal line = median and whiskers = min/max range). (B) A PCoA plot based on Bray–Curtis distance matrix. (C) Distribution of bacterial families between NABX and ABX. The composition of each family with relative abundance of more than 3 percent were illustrated. The parenthesis indicated phylum belonged to the family. (**p* < 0.05, ***p*< 0.01, ****p* < 0.001)

**Figure S11. Associations between microbiome and proteins in nasal secretions**

Spearman correlation heatmaps of total nasal microbiome (columns) and secreted proteome (row) in NABX (A) and ABX (B). The microbiome and proteome were arranged from top to down and from left to right, respectively, in order of the highest to the lowest number of significant correlations with each other. Black boxes indicated the top 25 percent microbiome and proteome with high number of significant correlations each other.

**Supplementary tables**

**Table S1. Identification of differentially expressed proteins with fold change ≥ 2.0 and *p*-value < 0.05 between disease status**

| Accession number | | Gene name | Log_2_ (Fold change) | - Log (*P*-value) |
| --- | --- | --- | --- | --- |
| CRSsNP vs. Control | Q9NYQ8 | FAT2 | 3.02 | 1.55 |
|  | P04003 | C4BPA | 1.39 | 2.69 |
|  | A6PVU8 | TUFT1 | 1.38 | 1.42 |
|  | A0A087X0P6 | IGKV2D-29 | 1.37 | 3.29 |
|  | P01031 | C5 | 1.35 | 3.31 |
|  | P23142 | FBLN1 | 1.33 | 4.69 |
|  | A0A087WSX0 | IGLV5-45 | 1.31 | 1.94 |
|  | P02747 | C1QC | 1.29 | 1.59 |
|  | P01616 |  | 1.29 | 2.10 |
|  | P08236 | GUSB | 1.13 | 1.61 |
|  | Q6P4A8 | PLBD1 | 1.12 | 1.36 |
|  | P01825 |  | 1.03 | 2.03 |
|  | A0A087X0N5 | IGKV1-17 | 1.02 | 2.06 |
|  | Q96QK1 | VPS35 | 1.01 | 1.37 |
|  | P68871 | HBB | -2.57 | 2.09 |
|  | G3V1N2 | HBA2 | -2.39 | 1.41 |
|  | P02042 | HBD | -2.30 | 1.91 |
|  | P69905 | HBA1 | -2.28 | 1.79 |
|  | P00915 | CA1 | -2.24 | 1.68 |
|  | P69891 | HBG1 | -2.00 | 1.74 |
|  | O15195 | VILL | -1.61 | 1.33 |
|  | P62328 | TMSB4X | -1.11 | 1.90 |
|  | P81605 | DCD | -1.11 | 2.33 |
|  | P00918 | CA2 | -1.08 | 2.07 |
|  | E9PN89 | HSPA8 | -1.05 | 1.35 |
| CRSwNP vs. Control | Q9NYQ8 | FAT2 | 3.43 | 1.80 |
|  | H3BT29 | PML | 1.98 | 1.61 |
|  | A0A075B6K3 | IGLV2-11 | 1.97 | 2.45 |
|  | A0A087WVM2 | CD177 | 1.89 | 2.23 |
|  | Q8WXI7 | MUC16 | 1.83 | 3.06 |
|  | Q86UN6 | AKAP14 | 1.71 | 2.03 |
|  | A0A075B7D0 | IGHV1OR15-1 | 1.68 | 2.01 |
|  | Q6UX06 | OLFM4 | 1.66 | 1.67 |
|  | P36222 | CHI3L1 | 1.51 | 1.80 |
|  | C9JC71 | FCGR3A | 1.49 | 1.45 |
|  | P01773 |  | 1.46 | 3.57 |
|  | Q92743 | HTRA1 | 1.46 | 1.70 |
|  | P01615 |  | 1.45 | 1.71 |
|  | A0A075B6H9 | IGLV4-69 | 1.44 | 1.34 |
|  | A0A075B6R9 | IGKV2D-24 | 1.43 | 3.01 |
|  | P01708 |  | 1.43 | 2.39 |
|  | P59665 | DEFA1 | 1.38 | 2.20 |
|  | P13671 | C6 | 1.36 | 1.84 |
|  | A0A087X0N5 | IGKV1-17 | 1.31 | 3.22 |
|  | O75884 | RBBP9 | 1.28 | 2.06 |
|  | A0A087X0P6 | IGKV2D-29 | 1.28 | 2.49 |
|  | A0A075B7B8 | IGHV3OR16-12 | 1.28 | 3.86 |
|  | Q5T5Y3 | CAMSAP1 | 1.26 | 1.66 |
|  | A0A096LPK4 | MUC5AC | 1.25 | 2.72 |
|  | Q15782 | CHI3L2 | 1.24 | 1.50 |
|  | P15328 | FOLR1 | 1.24 | 1.70 |
|  | P29401 | TKT | 1.23 | 2.51 |
|  | P05089 | ARG1 | 1.22 | 1.62 |
|  | P01707 |  | 1.19 | 1.67 |
|  | Q07654 | TFF3 | 1.15 | 1.97 |
|  | Q9UBG3 | CRNN | 1.14 | 2.03 |
|  | P01825 |  | 1.14 | 2.37 |
|  | O75348 | ATP6V1G1 | 1.12 | 2.18 |
|  | P49788 | RARRES1 | 1.11 | 1.42 |
|  | P08236 | GUSB | 1.10 | 1.55 |
|  | A0A087X0S5 | COL6A1 | 1.08 | 1.67 |
|  | P01031 | C5 | 1.08 | 2.18 |
|  | Q6ZVX7 | NCCRP1 | 1.07 | 1.62 |
|  | P23142 | FBLN1 | 1.06 | 3.05 |
|  | Q9BZG9 | LYNX1 | 1.06 | 2.83 |
|  | P61006 | RAB8A | 1.03 | 2.52 |
|  | P04209 |  | 1.01 | 1.46 |
|  | A0A075B6S3 | IGKV2-30 | 1.01 | 1.63 |
|  | O14773 | TPP1 | 1.01 | 1.39 |
|  | B1AKG0 | CFHR1 | 1.01 | 1.53 |
|  | Q02383 | SEMG2 | 1.00 | 1.46 |
|  | P04745 | AMY1A | 1.00 | 2.27 |
|  | G3V1N2 | HBA2 | -3.30 | 2.37 |
|  | P69905 | HBA1 | -2.78 | 2.51 |
|  | P69891 | HBG1 | -2.60 | 2.31 |
|  | P68871 | HBB | -2.56 | 2.38 |
|  | P00915 | CA1 | -2.54 | 1.96 |
|  | P02042 | HBD | -2.51 | 2.20 |
|  | P09210 | GSTA2 | -1.91 | 1.61 |
|  | Q6UWW0 | LCN15 | -1.87 | 1.66 |
|  | P11684 | SCGB1A1 | -1.79 | 4.03 |
|  | O15195 | VILL | -1.67 | 2.75 |
|  | A0A096LPE2 | SAA2-SAA4 | -1.67 | 1.92 |
|  | P42331-6 | ARHGAP25 | -1.67 | 2.65 |
|  | E9PN89 | HSPA8 | -1.52 | 2.86 |
|  | P16050 | ALOX15 | -1.52 | 1.46 |
|  | P19338 | NCL | -1.47 | 2.42 |
|  | P62857 | RPS28 | -1.40 | 2.11 |
|  | Q96KN2 | CNDP1 | -1.38 | 1.87 |
|  | C9J0K6 | SRI | -1.38 | 1.74 |
|  | P40394 | ADH7 | -1.33 | 2.24 |
|  | F6WQW2 | RANBP1 | -1.32 | 2.96 |
|  | Q96C23 | GALM | -1.29 | 2.18 |
|  | P02647 | APOA1 | -1.25 | 2.68 |
|  | J3QL71 | SCRN2 | -1.22 | 2.05 |
|  | P00326 | ADH1C | -1.21 | 1.52 |
|  | Q13442 | PDAP1 | -1.19 | 1.76 |
|  | Q13228 | SELENBP1 | -1.17 | 2.04 |
|  | P02768 | ALB | -1.16 | 2.39 |
|  | P02652 | APOA2 | -1.15 | 2.05 |
|  | P00918 | CA2 | -1.13 | 2.16 |
|  | A6NGP5 | HN1L | -1.10 | 1.66 |
|  | Q9H477 | RBKS | -1.09 | 1.53 |
|  | Q13885 | TUBB2A | -1.09 | 1.53 |
|  | P02794 | FTH1 | -1.07 | 1.35 |
|  | P61956 | SUMO2 | -1.06 | 2.44 |
|  | Q9H0E9 | BRD8 | -1.06 | 1.77 |
|  | M0R0K9 | TRIM28 | -1.03 | 1.62 |
|  | P09382 | LGALS1 | -1.02 | 3.26 |
|  | B8ZZQ6 | PTMA | -1.01 | 1.51 |
| CRSwNP vs. CRSsNP | C9JZR7 | ACTB | 2.08 | 1.92 |
|  | Q86UN6 | AKAP14 | 1.76 | 2.27 |
|  | Q04695 | KRT17 | 1.51 | 2.09 |
|  | B4DDF4 | CNN2 | 1.45 | 2.85 |
|  | Q9UBG3 | CRNN | 1.45 | 2.07 |
|  | P20930 | FLG | 1.40 | 1.57 |
|  | Q86UX7 | FERMT3 | 1.39 | 2.19 |
|  | Q7Z5R6 | APBB1IP | 1.36 | 1.97 |
|  | P05089 | ARG1 | 1.28 | 1.34 |
|  | Q8WXI7 | MUC16 | 1.27 | 1.66 |
|  | A0A087X188 | BIN2 | 1.22 | 1.78 |
|  | Q92743 | HTRA1 | 1.21 | 2.05 |
|  | Q9Y376 | CAB39 | 1.20 | 1.48 |
|  | Q9NUQ9 | FAM49B | 1.19 | 2.23 |
|  | A0A075B6R9 | IGKV2D-24 | 1.19 | 2.37 |
|  | P52790 | HK3 | 1.18 | 1.59 |
|  | Q9Y678 | COPG1 | 1.17 | 2.32 |
|  | P84085 | ARF5 | 1.14 | 2.63 |
|  | Q6UWP8 | SBSN | 1.14 | 1.89 |
|  | P29401 | TKT | 1.11 | 2.68 |
|  | O00203 | AP3B1 | 1.08 | 2.00 |
|  | P19878 | NCF2 | 1.08 | 1.59 |
|  | Q9Y490 | TLN1 | 1.06 | 2.20 |
|  | P26583 | HMGB2 | 1.06 | 2.05 |
|  | O75348 | ATP6V1G1 | 1.04 | 2.66 |
|  | Q13838 | DDX39B | 1.04 | 1.41 |
|  | P10644 | PRKAR1A | 1.02 | 1.56 |
|  | A0A087WV46 | LAMTOR4 | 1.02 | 1.40 |
|  | Q9HC84 | MUC5B | 1.01 | 1.61 |
|  | B1AH77 | RAC2 | 1.01 | 1.49 |
|  | P09210 | GSTA2 | -2.28 | 2.71 |
|  | A6PVU8 | TUFT1 | -2.03 | 3.17 |
|  | Q13938 | CAPS | -2.03 | 2.26 |
|  | Q96C23 | GALM | -1.64 | 2.77 |
|  | Q13885 | TUBB2A | -1.55 | 2.54 |
|  | C9J0K6 | SRI | -1.53 | 3.24 |
|  | P04003 | C4BPA | -1.49 | 3.01 |
|  | P02747 | C1QC | -1.44 | 2.26 |
|  | P00326 | ADH1C | -1.41 | 3.00 |
|  | A0A096LPE2 | SAA2-SAA4 | -1.40 | 1.59 |
|  | P62857 | RPS28 | -1.39 | 1.62 |
|  | P08263 | GSTA1 | -1.37 | 1.95 |
|  | Q9BW30 | TPPP3 | -1.37 | 1.84 |
|  | Q9H477 | RBKS | -1.36 | 2.45 |
|  | P51857 | AKR1D1 | -1.33 | 1.59 |
|  | P40394 | ADH7 | -1.29 | 2.28 |
|  | P11117 | ACP2 | -1.26 | 2.41 |
|  | P06576 | ATP5B | -1.25 | 3.17 |
|  | P12277 | CKB | -1.23 | 1.78 |
|  | P48595 | SERPINB10 | -1.18 | 1.71 |
|  | Q8WVM8 | SCFD1 | -1.11 | 1.49 |
|  | Q01105 | SET | -1.06 | 2.27 |
|  | Q99733 | NAP1L4 | -1.06 | 2.24 |
|  | K7EIJ8 | KATNAL2 | -1.04 | 2.19 |
|  | P23141 | CES1 | -1.04 | 1.62 |
|  | A6NGP5 | HN1L | -1.04 | 1.53 |
|  | Q8TD06 | AGR3 | -1.03 | 1.60 |
|  | Q13740 | ALCAM | -1.02 | 3.20 |

**Table S2. Identification of differentially expressed proteins with fold change ≥ 2.0 and *p*-value < 0.05 between Non-NP and NP in NABX**

| Accession number | | Gene name | Log_2_ (Fold change) | - Log (*P*-value) |
| --- | --- | --- | --- | --- |
| NP vs. Non-NP | Q9NYQ8 | FAT2 | 3.87 | 1.72 |
|  | A8MTF8 | FAM3B | 2.86 | 2.93 |
|  | Q9Y230 | RUVBL2 | 2.74 | 2.78 |
|  | C9JZR7 | ACTB | 2.52 | 2.36 |
|  | H3BT29 | PML | 2.22 | 1.59 |
|  | A0A087WVM2 | CD177 | 2.21 | 2.08 |
|  | P15814 | IGLL1 | 2.06 | 1.46 |
|  | A0A075B6R9 | IGKV2D-24 | 1.92 | 3.51 |
|  | Q9GZZ8 | LACRT | 1.87 | 2.07 |
|  | Q86UN6 | AKAP14 | 1.84 | 1.55 |
|  | P61626 | LYZ | 1.78 | 2.08 |
|  | A0A096LPK4 | MUC5AC | 1.75 | 3.83 |
|  | P47897 | QARS | 1.70 | 2.86 |
|  | P04206 |  | 1.61 | 1.60 |
|  | P01778 |  | 1.61 | 1.82 |
|  | Q96S96 | PEBP4 | 1.59 | 1.54 |
|  | P01708 |  | 1.57 | 2.49 |
|  | Q14240-2 | EIF4A2 | 1.53 | 1.64 |
|  | Q92882 | OSTF1 | 1.51 | 1.42 |
|  | Q6UX06 | OLFM4 | 1.50 | 1.49 |
|  | P18065 | IGFBP2 | 1.48 | 1.62 |
|  | Q6P5S2 | C6orf58 | 1.47 | 1.69 |
|  | A0A075B7D0 | IGHV1OR15-1 | 1.44 | 1.46 |
|  | P20827 | EFNA1 | 1.43 | 1.95 |
|  | Q9UBC9 | SPRR3 | 1.43 | 2.51 |
|  | O75884 | RBBP9 | 1.42 | 1.69 |
|  | P15328 | FOLR1 | 1.41 | 1.56 |
|  | Q5T5Y3 | CAMSAP1 | 1.41 | 1.83 |
|  | Q07654 | TFF3 | 1.40 | 2.04 |
|  | Q9HC84 | MUC5B | 1.40 | 1.78 |
|  | P29401 | TKT | 1.39 | 3.01 |
|  | Q9Y678 | COPG1 | 1.38 | 1.97 |
|  | A0A087WV46 | LAMTOR4 | 1.37 | 2.09 |
|  | Q8WXI7 | MUC16 | 1.37 | 1.35 |
|  | A0A087WZB2 | TYW1B | 1.37 | 1.39 |
|  | P49788 | RARRES1 | 1.31 | 1.41 |
|  | Q14204 | DYNC1H1 | 1.31 | 1.63 |
|  | Q86SQ4 | GPR126 | 1.30 | 1.76 |
|  | P00387-3 | CYB5R3 | 1.29 | 2.30 |
|  | J3KNB4 | CAMP | 1.29 | 1.34 |
|  | D6REX3 | SEC31A | 1.23 | 1.91 |
|  | P01615 |  | 1.21 | 1.57 |
|  | Q9UBG3 | CRNN | 1.19 | 1.35 |
|  | A0A087WXT3 | ZNF33B | 1.15 | 2.52 |
|  | P84085 | ARF5 | 1.12 | 2.36 |
|  | Q9BRX2 | PELO | 1.09 | 1.32 |
|  | P80188 | LCN2 | 1.09 | 3.69 |
|  | P01833 | PIGR | 1.09 | 1.89 |
|  | P09417 | QDPR | 1.07 | 1.34 |
|  | H0YL18 | B2M | 1.06 | 2.24 |
|  | A0A075B6S8 | IGKV1-5 | 1.06 | 5.17 |
|  | A0A087X0N5 | IGKV1-17 | 1.06 | 1.57 |
|  | P01621 |  | 1.05 | 3.43 |
|  | P78324 | SIRPA | 1.02 | 1.55 |
|  | P01773 |  | 1.00 | 1.90 |
|  | A0A096LPE2 | SAA2-SAA4 | -2.88 | 4.51 |
|  | B0YIW2 | APOC3 | -2.44 | 2.37 |
|  | P69905 | HBA1 | -2.35 | 1.38 |
|  | P69891 | HBG1 | -2.33 | 1.51 |
|  | P42331-6 | ARHGAP25 | -2.27 | 3.52 |
|  | E7ETZ0 | CALM1 | -2.14 | 1.56 |
|  | P34931 | HSPA1L | -2.13 | 1.42 |
|  | G3V2U4 | UNC79 | -2.11 | 2.03 |
|  | K7ER74 | APOC4-APOC2 | -2.06 | 2.11 |
|  | P33151 | CDH5 | -2.02 | 2.61 |
|  | P22891 | PROZ | -1.86 | 2.36 |
|  | P19338 | NCL | -1.83 | 2.21 |
|  | K7ERI9 | APOC1 | -1.83 | 2.33 |
|  | C9J0K6 | SRI | -1.79 | 2.38 |
|  | O75636 | FCN3 | -1.79 | 2.37 |
|  | Q96KN2 | CNDP1 | -1.75 | 2.90 |
|  | Q99733 | NAP1L4 | -1.73 | 4.38 |
|  | P02649 | APOE | -1.71 | 2.97 |
|  | O14791 | APOL1 | -1.70 | 1.80 |
|  | P04114 | APOB | -1.70 | 1.47 |
|  | Q6UWW0 | LCN15 | -1.70 | 1.33 |
|  | Q9H477 | RBKS | -1.69 | 2.33 |
|  | G3V0E5 | TFRC | -1.66 | 1.71 |
|  | P11684 | SCGB1A1 | -1.65 | 1.70 |
|  | Q6P387 | C16orf46 | -1.65 | 2.81 |
|  | P27169 | PON1 | -1.65 | 3.36 |
|  | P02747 | C1QC | -1.62 | 1.71 |
|  | P62701 | RPS4X | -1.61 | 3.38 |
|  | Q13885 | TUBB2A | -1.61 | 2.80 |
|  | H0Y8X4 | DNPH1 | -1.58 | 1.46 |
|  | E9PIA8 | PPT1 | -1.58 | 2.55 |
|  | P02652 | APOA2 | -1.55 | 3.25 |
|  | P23141 | CES1 | -1.52 | 1.85 |
|  | P02647 | APOA1 | -1.49 | 3.25 |
|  | P80108 | GPLD1 | -1.41 | 1.61 |
|  | P62857 | RPS28 | -1.40 | 1.78 |
|  | Q05BV3 | EML5 | -1.39 | 1.90 |
|  | Q15056 | EIF4H | -1.38 | 2.85 |
|  | G3XAL9 | SLC12A2 | -1.38 | 1.64 |
|  | P16050 | ALOX15 | -1.37 | 1.40 |
|  | Q9H4G0-2 | EPB41L1 | -1.34 | 2.01 |
|  | P52597 | HNRNPF | -1.34 | 2.47 |
|  | Q96C23 | GALM | -1.33 | 2.14 |
|  | P04180 | LCAT | -1.32 | 1.69 |
|  | A6NNI4 | CD9 | -1.30 | 1.81 |
|  | Q9UBE0 | SAE1 | -1.28 | 1.74 |
|  | P02794 | FTH1 | -1.27 | 1.64 |
|  | Q8WVM8 | SCFD1 | -1.24 | 1.88 |
|  | O95445 | APOM | -1.23 | 2.19 |
|  | Q15046 | KARS | -1.23 | 2.53 |
|  | P62805 | HIST1H4A | -1.23 | 1.34 |
|  | D6R967 | PPA2 | -1.21 | 2.83 |
|  | P05455 | SSB | -1.19 | 3.78 |
|  | A2A274 | ACO2 | -1.19 | 1.86 |
|  | Q14126 | DSG2 | -1.18 | 1.55 |
|  | P06576 | ATP5B | -1.15 | 2.53 |
|  | P11117 | ACP2 | -1.15 | 2.28 |
|  | P04003 | C4BPA | -1.15 | 1.40 |
|  | P01023 | A2M | -1.14 | 2.32 |
|  | A6NGP5 | HN1L | -1.14 | 1.57 |
|  | P09622 | DLD | -1.14 | 1.50 |
|  | Q15257 | PPP2R4 | -1.13 | 2.64 |
|  | P00326 | ADH1C | -1.12 | 1.37 |
|  | O94760 | DDAH1 | -1.12 | 1.76 |
|  | P98160 | HSPG2 | -1.11 | 1.68 |
|  | Q8WZA0 | LZIC | -1.10 | 2.50 |
|  | Q9H0E9 | BRD8 | -1.10 | 2.02 |
|  | O00410 | IPO5 | -1.10 | 1.56 |
|  | O43175 | PHGDH | -1.09 | 1.30 |
|  | P68371 | TUBB4B | -1.09 | 2.65 |
|  | Q15121 | PEA15 | -1.09 | 2.37 |
|  | Q96NY7 | CLIC6 | -1.09 | 2.05 |
|  | P12277 | CKB | -1.08 | 1.54 |
|  | P22792 | CPN2 | -1.07 | 2.11 |
|  | B7WNR0 | ALB | -1.07 | 3.07 |
|  | J3QL71 | SCRN2 | -1.07 | 1.30 |
|  | P00734 | F2 | -1.07 | 2.72 |
|  | D6W5Y5 | CIRBP | -1.06 | 1.42 |
|  | H3BM11 | SLC6A2 | -1.06 | 3.32 |
|  | P06727 | APOA4 | -1.06 | 2.53 |
|  | Q9H977 | WDR54 | -1.05 | 1.76 |
|  | Q9BX68 | HINT2 | -1.05 | 1.66 |
|  | Q9NQ48 | LZTFL1 | -1.04 | 1.70 |
|  | J3KN67 | TPM3 | -1.02 | 1.96 |
|  | Q9BW04 | SARG | -1.02 | 1.53 |
|  | Q03154 | ACY1 | -1.01 | 2.18 |
|  | F5GX07 | REXO2 | -1.01 | 2.73 |
|  | P07355 | ANXA2 | -1.01 | 2.71 |
|  | C9JF17 | APOD | -1.00 | 3.03 |

**Table S3. Identification of differentially expressed proteins with fold change ≥ 2.0 and *p*-value < 0.05 between Non-NP and NP in ABX**

| Accession number | | Gene name | Log_2_ (Fold change) | - Log (*P*-value) |
| --- | --- | --- | --- | --- |
| NP vs. Non-NP | Q86UN6 | AKAP14 | 3.39 | 3.28 |
|  | P33151 | CDH5 | 2.31 | 2.60 |
|  | Q9UGM5 | FETUB | 2.28 | 1.99 |
|  | P20930 | FLG | 2.08 | 1.46 |
|  | P13671 | C6 | 2.00 | 1.42 |
|  | Q6UWP8 | SBSN | 1.89 | 1.42 |
|  | E9PBS1 | PAICS | 1.88 | 2.01 |
|  | A6NLN1 | PTBP1 | 1.76 | 2.26 |
|  | P04180 | LCAT | 1.74 | 1.44 |
|  | P62263 | RPS14 | 1.71 | 1.66 |
|  | Q9BZG9 | LYNX1 | 1.66 | 2.18 |
|  | P00748 | F12 | 1.66 | 1.53 |
|  | O43390 | HNRNPR | 1.39 | 1.63 |
|  | P05198 | EIF2S1 | 1.36 | 1.50 |
|  | E7ETH6 | ZNF587B | 1.23 | 1.79 |
|  | P02753 | RBP4 | 1.15 | 1.73 |
|  | P10643 | C7 | 1.15 | 1.33 |
|  | E9PEB5 | FUBP1 | 1.13 | 1.70 |
|  | P01019 | AGT | 1.00 | 1.32 |
|  | Q9NYQ8 | FAT2 | -4.34 | 1.34 |
|  | E7EVA0 | MAP4 | -2.86 | 2.96 |
|  | P51857 | AKR1D1 | -2.54 | 1.45 |
|  | A0A087WW55 | PRSS1 | -2.53 | 1.33 |
|  | O76003 | GLRX3 | -2.23 | 1.84 |
|  | P48595 | SERPINB10 | -2.16 | 2.06 |
|  | O00757 | FBP2 | -1.78 | 1.44 |
|  | P52895 | AKR1C2 | -1.72 | 1.70 |
|  | Q04828 | AKR1C1 | -1.70 | 1.75 |
|  | P17050 | NAGA | -1.32 | 1.70 |
|  | Q9BRP8 | WIBG | -1.14 | 1.33 |

**Table S4. Identification of differentially expressed proteins with fold change ≥ 2.0 and *p*-value < 0.05 between NABX and ABX in total**

| Accession number | | Gene name | Log_2_ (Fold change) | - Log (*P*-value) |
| --- | --- | --- | --- | --- |
| ABX vs. NABX | A0A075B6H9 | IGLV4-69 | 2.14 | 2.37 |
|  | P00742 | F10 | 1.91 | 3.50 |
|  | Q9UN86 | G3BP2 | 1.76 | 5.91 |
|  | P08779 | KRT16 | 1.68 | 1.95 |
|  | P36222 | CHI3L1 | 1.59 | 2.26 |
|  | P05160 | F13B | 1.51 | 2.14 |
|  | E7END6 | PROC | 1.51 | 2.26 |
|  | Q7Z5R6 | APBB1IP | 1.49 | 3.00 |
|  | P20930 | FLG | 1.34 | 1.47 |
|  | Q6P4A8 | PLBD1 | 1.33 | 1.87 |
|  | P11234 | RALB | 1.29 | 2.49 |
|  | B7ZKW8 | RCSD1 | 1.26 | 3.30 |
|  | B4DUR8 | CCT3 | 1.20 | 1.58 |
|  | H0YLA2 | SRP14 | 1.18 | 3.36 |
|  | B1AKG0 | CFHR1 | 1.16 | 2.30 |
|  | P07948-2 | LYN | 1.15 | 1.79 |
|  | Q9Y376 | CAB39 | 1.13 | 1.67 |
|  | P80217 | IFI35 | 1.13 | 1.75 |
|  | Q8WXI7 | MUC16 | 1.11 | 1.35 |
|  | P78371 | CCT2 | 1.09 | 1.48 |
|  | P18428 | LBP | 1.09 | 1.57 |
|  | E9PG40 | APP | 1.08 | 1.85 |
|  | P29350 | PTPN6 | 1.08 | 1.59 |
|  | C9JB55 | TF | 1.06 | 1.35 |
|  | Q86YZ3 | HRNR | 1.03 | 2.07 |
|  | P27361 | MAPK3 | 1.02 | 1.53 |
|  | P13647 | KRT5 | 1.01 | 2.35 |
|  | Q5THJ4 | VPS13D | -2.55 | 2.63 |
|  | O14948 | TFEC | -2.37 | 1.89 |
|  | O15195 | VILL | -2.23 | 2.56 |
|  | F6TR53 | HS1BP3 | -1.78 | 3.01 |
|  | P47929 | LGALS7 | -1.70 | 2.14 |
|  | H0YLI6 | IDH3A | -1.68 | 2.96 |
|  | A0A087WZW8 | IGKV3-11 | -1.65 | 2.85 |
|  | P21964 | COMT | -1.42 | 1.87 |
|  | P01742 |  | -1.38 | 2.23 |
|  | Q14258 | TRIM25 | -1.38 | 1.51 |
|  | A0A087WZG4 | ARHGEF18 | -1.37 | 1.50 |
|  | P01703 |  | -1.33 | 2.57 |
|  | P01778 |  | -1.32 | 1.52 |
|  | P62328 | TMSB4X | -1.31 | 2.25 |
|  | P01699 |  | -1.22 | 1.98 |
|  | P63313 | TMSB10 | -1.20 | 3.29 |
|  | P01772 |  | -1.18 | 2.71 |
|  | P40394 | ADH7 | -1.17 | 2.10 |
|  | P62857 | RPS28 | -1.17 | 1.61 |
|  | P01700 |  | -1.17 | 1.74 |
|  | P01763 |  | -1.17 | 2.51 |
|  | F8VXU5 | VPS29 | -1.14 | 2.44 |
|  | F8VVT9 | AGAP2 | -1.14 | 2.46 |
|  | H7BZJ3 | PDIA3 | -1.13 | 2.55 |
|  | P04792 | HSPB1 | -1.10 | 2.80 |
|  | K7EJC1 | PSMD8 | -1.10 | 1.64 |
|  | F6WQW2 | RANBP1 | -1.09 | 3.44 |
|  | Q96C23 | GALM | -1.09 | 1.87 |
|  | B8ZZQ6 | PTMA | -1.06 | 1.70 |
|  | P16403 | HIST1H1C | -1.05 | 1.46 |
|  | P01876 | IGHA1 | -1.02 | 3.70 |
|  | O95881 | TXNDC12 | -1.01 | 4.76 |
|  | P06703 | S100A6 | -1.01 | 2.32 |

**Table S5. Identification of differentially expressed proteins with fold change ≥ 2.0 and *p*-value < 0.05 between NABX and ABX in Non-NP**

| Accession number | | Gene name | Log_2_ (Fold change) | - Log (*P*-value) |
| --- | --- | --- | --- | --- |
| ABX vs. NABX | Q9NYQ8 | FAT2 | 4.06 | 1.31 |
|  | A0A087WW55 | PRSS1 | 2.49 | 1.50 |
|  | O75976 | CPD | 2.24 | 2.45 |
|  | E7END6 | PROC | 1.75 | 1.45 |
|  | A0A075B6J3 | IGLV3-27 | 1.68 | 2.39 |
|  | B4DUR8 | CCT3 | 1.62 | 1.44 |
|  | E7EVA0 | MAP4 | 1.56 | 1.43 |
|  | Q16719 | KYNU | 1.52 | 2.05 |
|  | A0A087WUQ6 | GPX1 | 1.45 | 1.54 |
|  | P01781 |  | 1.42 | 2.01 |
|  | Q9BRX2 | PELO | 1.39 | 1.51 |
|  | P12429 | ANXA3 | 1.29 | 1.33 |
|  | Q12765 | SCRN1 | 1.29 | 1.99 |
|  | Q9UHJ6 | SHPK | 1.27 | 2.55 |
|  | P48739 | PITPNB | 1.27 | 1.63 |
|  | Q13185 | CBX3 | 1.20 | 1.86 |
|  | P53634 | CTSC | 1.19 | 2.94 |
|  | B7ZKW8 | RCSD1 | 1.14 | 1.35 |
|  | O00754 | MAN2B1 | 1.11 | 1.61 |
|  | P11279 | LAMP1 | 1.10 | 1.35 |
|  | Q8WW12 | PCNP | 1.05 | 1.90 |
|  | Q9BVG4 | PBDC1 | 1.00 | 1.33 |
|  | O15195 | VILL | -3.65 | 2.69 |
|  | F6TR53 | HS1BP3 | -2.59 | 2.82 |
|  | K7ER74 | APOC4-APOC2 | -2.45 | 1.86 |
|  | A0A087WZW8 | IGKV3-11 | -2.12 | 2.26 |
|  | H0YLI6 | IDH3A | -2.01 | 2.58 |
|  | G3V0E5 | TFRC | -1.96 | 1.41 |
|  | P16403 | HIST1H1C | -1.73 | 1.75 |
|  | P01772 |  | -1.70 | 2.41 |
|  | P01699 |  | -1.68 | 1.58 |
|  | E7EU04 | FOLR2 | -1.66 | 2.28 |
|  | P63313 | TMSB10 | -1.66 | 2.41 |
|  | Q9BW04 | SARG | -1.60 | 2.09 |
|  | P01703 |  | -1.54 | 1.65 |
|  | P01700 |  | -1.52 | 1.31 |
|  | A0A075B6F3 | DNAH8 | -1.52 | 2.72 |
|  | P62195 | PSMC5 | -1.52 | 1.68 |
|  | P02652 | APOA2 | -1.43 | 1.55 |
|  | F2Z2G4 | WFDC3 | -1.36 | 1.45 |
|  | P01608 |  | -1.21 | 1.70 |
|  | J3KP15 | SRSF2 | -1.20 | 1.90 |
|  | P16949 | STMN1 | -1.18 | 2.35 |
|  | P00966 | ASS1 | -1.15 | 2.22 |
|  | P09382 | LGALS1 | -1.14 | 1.83 |
|  | A0A075B6J9 | IGLV2-18 | -1.14 | 1.53 |
|  | E7ETH6 | ZNF587B | -1.12 | 2.30 |
|  | P62318 | SNRPD3 | -1.11 | 1.37 |
|  | A0A087WSY5 | CPB2 | -1.05 | 1.71 |
|  | P09238 | MMP10 | -1.05 | 2.35 |

**Table S6. Identification of differentially expressed proteins with fold change ≥ 2.0 and *p*-value < 0.05 between NABX and ABX in NP**

| Accession number | | Gene name | Log_2_ (Fold change) | - Log (*P*-value) |
| --- | --- | --- | --- | --- |
| ABX vs. NABX | P01775 |  | 4.33 | 1.68 |
|  | G3V2U4 | UNC79 | 3.34 | 2.62 |
|  | P33151 | CDH5 | 3.27 | 4.84 |
|  | C9JB55 | TF | 2.92 | 3.15 |
|  | A0A096LPE2 | SAA2-SAA4 | 2.80 | 2.24 |
|  | A0A075B6H9 | IGLV4-69 | 2.78 | 2.06 |
|  | P05160 | F13B | 2.58 | 1.91 |
|  | O14791 | APOL1 | 2.52 | 2.66 |
|  | B0YIW2 | APOC3 | 2.50 | 1.68 |
|  | Q9UN86 | G3BP2 | 2.44 | 3.11 |
|  | P80108 | GPLD1 | 2.43 | 2.51 |
|  | Q9P265 | DIP2B | 2.38 | 4.03 |
|  | O75636 | FCN3 | 2.36 | 3.04 |
|  | F8WBL1 | L3MBTL2 | 2.33 | 1.60 |
|  | A0A087WUS7 | IGHD | 2.32 | 1.57 |
|  | D6RAT0 | RPS3A | 2.31 | 1.66 |
|  | P62805 | HIST1H4A | 2.29 | 1.89 |
|  | H0YLA2 | SRP14 | 2.17 | 3.82 |
|  | P20930 | FLG | 2.14 | 2.00 |
|  | P00742 | F10 | 2.09 | 1.83 |
|  | P04180 | LCAT | 2.05 | 3.63 |
|  | Q05BV3 | EML5 | 2.04 | 1.49 |
|  | P09622 | DLD | 2.03 | 1.89 |
|  | P27169 | PON1 | 1.98 | 3.38 |
|  | Q96JB5 | CDK5RAP3 | 1.97 | 2.43 |
|  | P18428 | LBP | 1.97 | 1.72 |
|  | P31942 | HNRNPH3 | 1.97 | 1.54 |
|  | K7ERI9 | APOC1 | 1.94 | 1.48 |
|  | B1AKG0 | CFHR1 | 1.89 | 2.15 |
|  | P14923 | JUP | 1.87 | 1.92 |
|  | F8WF14 | BCHE | 1.82 | 1.55 |
|  | Q9Y570 | PPME1 | 1.80 | 1.97 |
|  | P04839 | CYBB | 1.79 | 1.35 |
|  | Q9UNH7 | SNX6 | 1.79 | 2.20 |
|  | P02649 | APOE | 1.78 | 2.09 |
|  | Q6UWP8-2 | SBSN | 1.73 | 1.64 |
|  | B7ZKW8 | RCSD1 | 1.69 | 1.75 |
|  | P11234 | RALB | 1.69 | 1.63 |
|  | P41250 | GARS | 1.68 | 1.93 |
|  | P80217 | IFI35 | 1.67 | 1.60 |
|  | Q9UGM5 | FETUB | 1.63 | 1.44 |
|  | Q9UBE0 | SAE1 | 1.63 | 1.56 |
|  | Q6P387 | C16orf46 | 1.62 | 1.59 |
|  | P13647 | KRT5 | 1.62 | 2.09 |
|  | P35908 | KRT2 | 1.57 | 1.58 |
|  | P19338 | NCL | 1.56 | 1.65 |
|  | Q8NFL0 | B3GNT7 | 1.55 | 1.38 |
|  | P34913 | EPHX2 | 1.54 | 2.62 |
|  | P00739 | HPR | 1.53 | 2.09 |
|  | P22792 | CPN2 | 1.51 | 1.93 |
|  | P07305 | H1F0 | 1.50 | 2.01 |
|  | P05154 | SERPINA5 | 1.49 | 2.60 |
|  | P02753 | RBP4 | 1.49 | 3.94 |
|  | Q9Y536 | PPIAL4A | 1.48 | 1.75 |
|  | Q9HC10 | OTOF | 1.47 | 2.24 |
|  | C9JXI5 | TMEM198 | 1.46 | 1.79 |
|  | P43652 | AFM | 1.46 | 2.07 |
|  | P05388 | RPLP0 | 1.46 | 1.85 |
|  | Q06033 | ITIH3 | 1.44 | 2.06 |
|  | O95445 | APOM | 1.42 | 1.61 |
|  | P05546 | SERPIND1 | 1.42 | 2.56 |
|  | Q86YZ3 | HRNR | 1.41 | 1.38 |
|  | Q8N1G4 | LRRC47 | 1.38 | 2.04 |
|  | O60814 | HIST1H2BK | 1.37 | 1.62 |
|  | F5H5U2 | DDX55 | 1.33 | 2.89 |
|  | C9JF17 | APOD | 1.32 | 3.05 |
|  | P15169 | CPN1 | 1.32 | 2.06 |
|  | Q9P1F3 | ABRACL | 1.31 | 1.74 |
|  | P01042-2 | KNG1 | 1.31 | 3.18 |
|  | P04217-2 | A1BG | 1.30 | 2.60 |
|  | P07358 | C8B | 1.30 | 1.59 |
|  | Q8NC51 | SERBP1 | 1.30 | 2.59 |
|  | Q5SSJ5 | HP1BP3 | 1.30 | 1.62 |
|  | P55884 | EIF3B | 1.28 | 1.90 |
|  | Q9H0P0-1 | NT5C3A | 1.26 | 1.68 |
|  | P00748 | F12 | 1.26 | 1.69 |
|  | Q14624 | ITIH4 | 1.24 | 2.40 |
|  | Q9P258 | RCC2 | 1.24 | 1.70 |
|  | E9PM52 | SIRT3 | 1.24 | 1.37 |
|  | A0A087X232 | C1S | 1.23 | 1.80 |
|  | Q8WW12 | PCNP | 1.22 | 1.45 |
|  | P12429 | ANXA3 | 1.22 | 1.73 |
|  | H3BM11 | SLC6A2 | 1.22 | 2.85 |
|  | Q12882 | DPYD | 1.21 | 1.94 |
|  | B4DEB1 | H3F3A | 1.21 | 1.56 |
|  | B5MBZ0 | EML4 | 1.20 | 2.04 |
|  | P07225 | PROS1 | 1.18 | 1.90 |
|  | D6R967 | PPA2 | 1.14 | 2.23 |
|  | J3KN67 | TPM3 | 1.13 | 1.51 |
|  | Q8WZA0 | LZIC | 1.13 | 1.45 |
|  | P00734 | F2 | 1.12 | 1.61 |
|  | P02743 | APCS | 1.11 | 2.25 |
|  | O00487 | PSMD14 | 1.11 | 1.70 |
|  | Q8TDW7 | FAT3 | 1.10 | 1.45 |
|  | P05114 | HMGN1 | 1.09 | 1.40 |
|  | P05198 | EIF2S1 | 1.08 | 1.40 |
|  | Q96HC4 | PDLIM5 | 1.07 | 1.35 |
|  | A0A087WVQ6 | CLTC | 1.07 | 2.02 |
|  | P55036 | PSMD4 | 1.05 | 1.46 |
|  | Q3YEC7 | RABL6 | 1.03 | 1.39 |
|  | Q9NYQ8 | FAT2 | -3.62 | 1.33 |
|  | Q5THJ4 | VPS13D | -2.95 | 1.62 |
|  | H3BT29 | PML | -2.35 | 1.67 |
|  | A0A087WZB2 | TYW1B | -2.35 | 1.54 |
|  | P47929 | LGALS7 | -2.31 | 1.45 |
|  | P15814 | IGLL1 | -2.22 | 1.72 |
|  | A0A087WZW8 | IGKV3-11 | -2.18 | 1.76 |
|  | P01778 |  | -2.15 | 1.94 |
|  | E7EVA0 | MAP4 | -2.14 | 1.73 |
|  | P01742 |  | -2.10 | 1.79 |
|  | P61626 | LYZ | -2.10 | 2.38 |
|  | P01833 | PIGR | -2.08 | 2.32 |
|  | P02788 | LTF | -1.99 | 1.89 |
|  | P01877 | IGHA2 | -1.91 | 2.60 |
|  | P01876 | IGHA1 | -1.86 | 2.41 |
|  | P01703 |  | -1.85 | 1.79 |
|  | Q6P5S2 | C6orf58 | -1.75 | 1.42 |
|  | K7EJC1 | PSMD8 | -1.69 | 1.80 |
|  | Q9UBC9 | SPRR3 | -1.69 | 1.88 |
|  | Q9UGM3 | DMBT1 | -1.62 | 2.02 |
|  | P00387-3 | CYB5R3 | -1.62 | 2.00 |
|  | P06703 | S100A6 | -1.59 | 2.28 |
|  | P25815 | S100P | -1.57 | 2.47 |
|  | Q14566 | MCM6 | -1.54 | 1.60 |
|  | Q01469 | FABP5 | -1.49 | 2.15 |
|  | H0YL18 | B2M | -1.49 | 2.04 |
|  | O00757 | FBP2 | -1.48 | 2.34 |
|  | P25311 | AZGP1 | -1.47 | 2.26 |
|  | P48594 | SERPINB4 | -1.44 | 1.39 |
|  | P31949 | S100A11 | -1.43 | 2.27 |
|  | F8VXU5 | VPS29 | -1.41 | 1.72 |
|  | P01772 |  | -1.40 | 1.56 |
|  | Q8TAX7 | MUC7 | -1.39 | 1.35 |
|  | Q9C005 | DPY30 | -1.38 | 1.49 |
|  | Q92616 | GCN1L1 | -1.38 | 1.62 |
|  | P18510-2 | IL1RN | -1.37 | 1.74 |
|  | F8VVT9 | AGAP2 | -1.34 | 1.77 |
|  | A0A087WYR4 | IGLL5 | -1.34 | 3.19 |
|  | P01708 |  | -1.32 | 1.45 |
|  | A0A075B7E8 | IGHV3OR16-13 | -1.30 | 1.43 |
|  | P01621 |  | -1.30 | 2.19 |
|  | P01591 | IGJ | -1.29 | 2.75 |
|  | P23528 | CFL1 | -1.28 | 1.69 |
|  | A0A075B6R9 | IGKV2D-24 | -1.27 | 2.42 |
|  | A0A075B6K4 | IGLV3-10 | -1.27 | 1.31 |
|  | P17931 | LGALS3 | -1.25 | 1.92 |
|  | P20061 | TCN1 | -1.25 | 1.85 |
|  | P06733 | ENO1 | -1.24 | 1.96 |
|  | A0A075B6K9 | IGLC2 | -1.23 | 2.22 |
|  | O95994 | AGR2 | -1.22 | 2.06 |
|  | P26447 | S100A4 | -1.21 | 2.27 |
|  | H7BZJ3 | PDIA3 | -1.20 | 1.59 |
|  | P01034 | CST3 | -1.20 | 1.75 |
|  | P17066 | HSPA6 | -1.16 | 1.64 |
|  | E7ES19 | THBS4 | -1.16 | 1.90 |
|  | P61960 | UFM1 | -1.15 | 1.36 |
|  | A0A096LPK4 | MUC5AC | -1.14 | 1.33 |
|  | P60709 | ACTB | -1.13 | 1.44 |
|  | P80188 | LCN2 | -1.12 | 1.80 |
|  | P04207 |  | -1.11 | 1.85 |
|  | A0A075B6S8 | IGKV1-5 | -1.09 | 2.62 |
|  | P01610 |  | -1.07 | 1.81 |
|  | P31947 | SFN | -1.05 | 2.32 |
|  | A0A087WYL9 | IGKC | -1.03 | 3.10 |
|  | A0A087X1V9 | IGKV2-28 | -1.02 | 2.73 |
|  | A0A087WV47 | IGHG1 | -1.01 | 1.73 |
